# Evolution of two gene networks underlying adaptation to drought stress in the wild tomato *Solanum chilense*

**DOI:** 10.1101/2023.01.18.524537

**Authors:** Kai Wei, Saida Sharifova, Xiaoyun Zhao, Neelima Sinha, Hokuto Nakayama, Aurélien Tellier, Gustavo A Silva-Arias

## Abstract

Drought stress is a key factor limiting plant growth and the colonization of arid habitats by plants. Here, we study the evolution of gene expression response to drought stress in a wild tomato, *Solanum chilense* naturally occurring around the Atacama Desert in South America. We conduct a transcriptome analysis of plants under standard and drought experimental conditions to understand the evolution of drought-response gene networks. We identify two main regulatory networks corresponding to two typical drought-responsive strategies: cell cycle and fundamental metabolic processes. We estimate the age of the genes in these networks and the age of the gene expression network, revealing that the metabolic network has a younger origin and more variable transcriptome than the cell-cycle network. Combining with analyses of population genetics, we found that a higher proportion of the metabolic network genes show signatures of recent positive selection underlying recent adaptation within *S. chilense,* while the cell-cycle network appears of ancient origin and is more conserved. For both networks, however, we find that genes showing older age of selective sweeps are the more connected in the network. Adaptation to southern arid habitats over the last 50,000 years occurred in *S. chilense* by adaptive changes core genes with substantial network rewiring and subsequently by smaller changes at peripheral genes.

## Introduction

Drought stress is one of the major environmental constraints negatively influencing plant development and preventing plant growth, resulting in decreased yield in agriculture and as a constraining factor for colonization of arid or hyper-arid habitats (Ciais et al. 2005; Juenger 2013). Plants respond to water-insufficiency through multiple strategies underpinned by various physiological and developmental processes, such as storage of internal water to avoid tissue damage and tolerance (endurance) to drought stress to maintain the growth process (Basu et al. 2016). These strategies involve many biological functions such as increasing the metabolic activity of some tissues, i.e. root water uptake and closing stomata, or activation of metabolic pathways including phytohormone signaling, antioxidant and metabolite production in order to regulate osmotic processes (Rodrigues et al. 2019). Drought response involves numerous quantitative and polygenic traits acting in (complex) gene co-expression networks (GCN). To improve crops and predict the evolutionary responses of plant species under the current and predicted global water deficits, it is thus of interest to pinpoint and decipher the evolutionary history of the relevant GCNs underpinning the adaptation of wild plants to arid or hyper-arid habitats (Gehan et al. 2015).

Comparative transcriptomics involving the inference of gene co-expression patterns show that many GCNs are conserved through the tree of life (Stuart et al. 2003; Gerstein et al. 2014; Zarrineh et al. 2014; Crow et al. 2022). Moreover, phylogenetic and developmental studies have demonstrated that many physiological, structural, and regulatory innovations to cope drought stress have arisen throughout the history of plants, many of them even predating the emergence of land plants (Jill Harrison 2017; de Vries et al. 2018; de Vries and Archibald 2018; Mustafin et al. 2019; Wang et al. 2020; Bowles et al. 2021). Several conserved GCNs can be observed in fundamental biological processes such as protein metabolism, cell cycle, and photosynthesis and well as key traits such wood formation (Stuart et al. 2003; Ficklin and Feltus 2011; Zinkgraf et al. 2020).

A key question in functional and evolutionary genomics is thus to link GCN evolution and (relatively) short-scale evolutionary processes such as adaptation and population/species divergence in order to assess the relative importance of contingency, exaptation and evolution of novel genes (duplication, neofunctionalization) allowing colonization of novel habitats. Two main hypotheses are formulated. First, highly conserved sub-networks (so-called hubs or kernels) evolve under strong purifying selection to ensure the functionality of the GCNs (Papakostas et al. 2014; Josephs et al. 2017; Mähler et al. 2017; Masalia et al. 2017), so that genetic variation is only found at (less connected) genes at the periphery of the GCNs that may be the target of positive natural selection (Flowers et al. 2007; Kim et al. 2007; Luisi et al. 2015; Erwin 2020). However, this argument is likely because the novel habitats may not differ much from the original one, so that only minor adjustments in the GCNs are enough to provide adaptation. This is also in line with so-called developmental systems drift (DSD; True and Haag 2001), that predicts GCN rewiring only occurs in ‘flexible’ (sub-)modules with the accumulation of neutral variation that keep the network function intact until a new viable function (phenotype or developmental pathway) appears. Second, despite the general belief that genes with higher connectivity evolve at a slower rate, there is also evidence that changes at central genes (with higher connectivity) can be responsible for the short-term response to selection (Jovelin and Phillips 2009; Luisi et al. 2015) and promote rewiring of the GCN (Koubkova-Yu et al. 2018). Thus, highly connected genes may be targets of positive selection during environmental change, *e.g.* adaptation to novel habitats, even though these genes experience purifying selection in stable environments (Hämälä et al. 2020). Indeed, if the second hypothesis is correct, we expect a correlation between the age of positive selection and the connectivity of a gene in a network, but no correlation under the first hypothesis.

To test these hypotheses, we reveal the selective forces (positive versus purifying selection) acting on different components of the networks (hub vs peripheral genes) across species/lineages adapted to contrasting conditions, and correlate the signals of positive selection with gene connectivity in the wild tomato species *Solanum chilense*. Wild tomatoes are a model of interest as their diversification is accompanied by the exploration of wide environmental gradients along the Pacific coast of South America (from tropical to subtropical, coastal to high mountain, and wet to extremely dry regions; Nakazato et al. 2010; Haak et al. 2014). In addition, the infra-specific diversification within *S. chilense* resulted in several lineages with strong environmental differentiation (Raduski and Igić 2021; Wei et al. 2023). Populations of *S. chilense* are challenged by prolonged drought, with the most severe drought conditions occurring in the southern part of the range. Wild relative tomato species such *S. chilense*, *S. sitiens* and *S. pennellii* become well-established systems to study tolerance strategies to survive in extreme environments (Bolger et al. 2014; Martínez et al. 2014; Tapia et al. 2016; Kashyap et al. 2020; Blanchard-Gros et al. 2021; Molitor et al. 2021; Barrera-Ayala et al. 2023). In a previous study, we assayed for evidences of positive selection in 30 fully sequenced genomes of *S. chilense* to identify candidate genes underpinning adaptation along the species range. We found genes with putative functions related to root hair development and cell homeostasis as being likely involved in drought stress tolerance (Wei et al. 2023). However, to date, most research in *S. chilense* has focused on the evolution of a few genes potentially involved in abiotic stress response (Fischer et al. 2011; Mboup et al. 2012; Fischer et al. 2013; Böndel et al. 2015; Nosenko et al. 2016; Böndel et al. 2018), and we still lack information regarding the evolutionary mechanisms driving drought tolerance in this species.

Our aim is to study the GCN evolution underpinning *S. chilense* adaptation to arid habitats. We identify drought stress responsive gene regulatory networks combining multiple analyses of transcriptome data of *S. chilense* and focus on two networks involved in cell-cycle and metabolic processes. Furthermore, we infer the evolutionary processes at these two networks across three different evolutionary levels (tree of life/plants, species and population) by computation of transcriptome indices to explore the evolutionary age and sequence divergence of the drought responsive transcriptome. We then analyze the emergence of adaptive variation in the identified drought-responsive genes of these networks and the association to gene connectivity.

## Results

### Drought experiments and transcriptome analyses

Plants of *S. chilense* growing either under well-watered or a moderate water stress regimes (hereafter, control and drought) showed clear morphological differences by the time of tissue collection. Plant growth and ramification was boosted in well-watered group while plants under drought were smaller and slow-growing. Hence, on the day 12, newly expanded leaf and shoot apices were collected for the expression analysis of stress-responsive genes and four biological replicates were used for all RNA-Seq experiments from each tissue type.

We analyze short-read transcriptome data from 16 libraries aligned to the reference genome of *S. chilense* (Dataset S1). A total of 27,832 genes are identified to be expressed in the 16 libraries (Dataset S2), of which 1,536 genes are uniquely expressed in drought condition and 1,767 genes in control condition (Dataset S2). A principal component analysis (PCA) based on the gene expression profiles reveals consistent clustering primarily associated with the experimental conditions (control and drought) and secondarily to the developmental stages (leaf and shoot apex) (Figure 1A). PC1 accounts for 28.17% of the expression variability and separates the libraries from the two experimental conditions, indicating transcriptome remodeling between drought and control conditions. Libraries from different developmental stages are separated along the PC2 axis (accounting for 18.24% of the variance), supporting tissue age transcriptome specificity. Consistently, the transcriptome similarity analysis between libraries reveals that the watering conditions explain the major differences between treatments (Figure 1B). Hierarchical clustering also revealed that the transcriptomes were grouped mainly according to water deficit intensity, rather than by tissue type (Figure 1C), demonstrating the predominant effect of the stress response in transcriptome remodeling. Therefore, we thereafter focus on comparing the transcriptome profiles of the drought and control experimental conditions.

### Identification of gene networks involved in drought stress

We identified gene networks involved in drought response in *S. chilense* based on differential expression analysis and weighted gene co-expression network analysis (WGCNA). First, three sets of differential expression genes (DEGs) are identified from three drought/control comparison groups (full data set, only leaf and only shoot apex tissues) (Figure 2A; Dataset S3; log2FoldChange ≥ 1, FDR *P* ≤ 0.001). A total of 4,905 DEGs are obtained in three comparison groups, of which 2,484 DEGs (1,235 up-regulated and 1,249 down-regulated in drought transcriptome) are shared in three comparison groups (Figure 2B). We deduce that these shared DEGs correspond to a core functionally drought-responsive network.

In construction of gene co-expression networks (GCNs), we do not directly used DEGs in WGCNA as suggested by the developer of WGCNA, because DEGs are invalid for assumption of the scale-free topology. Therefore, a set of 16,181 genes after filtering from all expressed genes were used in WGCNA (see methods), and clustered into seven co-expression modules named after different colors. The module sizes range from 183 up to 5,364 genes (Figure 2C, Dataset S4). Among the identified co-expression modules, the blue module (3,852 genes) shows significantly positive correlation with control condition and negative correlation with drought condition (Figure 2C, Kendall’s test, *P* = 2.2e-11). In contrast, the turquoise module (5,364 genes) is significantly positively correlated with drought condition and negatively correlated with control condition (Figure 2C, Kendall’s test, *P* = 2.34e-13). In addition, the genes within blue and turquoise modules are observed to show higher connectivity than other modules (Figure S1, Kolmogorov-Smirnov test on connectivity measure, *P* = 2.41e-17), indicating higher interaction and closer correspondence in biological process among genes within each module in response to water deprivation.

We next check the overlap between 2,484 DEGs and co-expression modules to confirm that blue and turquoise modules are associated with drought stress in *S. chilense* (Table S1). DEGs share far more genes with the blue and turquoise modules than with other co-expression modules. Almost all shared DEGs (2,302 genes out of 2,484) are found in the blue and turquoise modules. This confirms that blue and turquoise modules are two sets of co-expressed drought stress responsive genes. The overlapping DEGs and module genes are extracted to constitute the now refined two high-confidence subsets of the blue and turquoise modules and comprising 1,223 and 1,079 genes, respectively.

To independently support regulatory relationships among genes identified in the two co-expression networks, we identify transcription factors (TFs) and transcription factor binding sites (TFBSs) for the two subsets of genes. Therefore, we extract the genes that can bind to one another (Table S2) from the two high-confidence subsets, which we hereafter name as sub-blue (686 genes) and sub-turquoise (948 genes), respectively (Dataset S5). These results show that genes in the sub-blue and sub-turquoise networks not only show specific co-expression patterns, but they show also predicted interact between the TFs and TFBSs. Subsequently, the co-expression network is reconstructed using the same steps for the set of genes of the sub-blue and sub-turquoise networks. Higher connectivity is observed in the sub-turquoise network (Figure S2, Kolmogorov-Smirnov test on connectivity measure, *P =* 0.002), suggesting a closer regulatory relationship among genes in the sub-turquoise than in the sub-blue network.

To verify the robustness of two drought-responsive network (sub-blue and sub-turquoise) across different tomato species (and thus the generality of our results), we employed also the same pipeline to construct GCNs combined to transcriptomic data of *S. pennelliii* (PRJEB5809; Bolger et al. 2014) and *S. lycopersicum* (PRJNA812356; Yang et al. 2022) (Dataset S1). Those transcriptomes also exhibit difference due to water conditions in the analysis of PCA, correlations and hierarchical clustering (Figure S3). We first observe that the 1,837 (74%) genes in the DEG sets based on only *S. chilense* overlap to the combined DEG set of *S. pennellii* and *S*. *lycopersicum* (Figure S4A and S4B). The GCNs from combined transcriptome profiles consistently show two networks as we find in *S. chilense* alone (Figure 2C and S4C). We find that 576 (84%) and 778 (82%) genes overlap to the sub-blue and sub-turquoise networks, respectively (Figure S4D and S4E). In addition, our two drought-responsive networks (sub-blue and sub-turquoise) also overlap with previous study based on drought transcriptomes of *S. lycopersicum* (Nicolas et al. 2022). Nearly 60% of drought-responsive genes in the two networks are observed as response to water stress in *S. lycopersicum*. These overlap rates suggest that our two drought-responsive networks are present and perform similar functions in different tomato species. We therefore concentrate on these two networks: sub-blue (686 genes) and sub-turquoise (948 genes).

### Functional enrichment analysis of drought-responsive GCNs

We assess whether the two identified gene networks (sub-blue and sub-turquoise) show functional differences. The gene ontology (GO) enrichment reveals that sub-blue network is significantly enriched (*P* < 0.05) in cell cycle and regulation biological processes, including replication and modification of genetic information, ribosome production and assembly, cytoskeleton organization, among others (Figure 3A; Table S3). Conversely, the sub-turquoise network is enriched in biological processes related to response of physiological and metabolic processes to water shortage and heat, including some metabolic processes, signal pathways, changes of stomata and cuticle, amongst other processes (Figure 3A; Table S3). These functional differences suggest that genes in the two sub-networks are activated and expressed in different cellular compartments. Consistent with the mentioned biological process, the sub-blue network genes are mainly enriched in cellular components in the nucleus, including nucleolus, chromosome, nuclear envelope, and ribosome (Figure 3B; Table S4). These cellular components are at the center of cell division processes. On the other hand, the sub-turquoise network is enriched in cellular components related to metabolism processes, such as complexes and membrane structures in the cell (Figure 3B; Table S4). Many studies have indicated that modulation in the cell cycle and fundamental metabolism are two main strategies in response to drought stress (Gupta et al. 2020; Yang et al. 2021; Nicolas et al. 2022). We focus, thereafter, on these two sub-networks and from now on, the sub-blue network is referred to as the *cell-cycle network* and the sub-turquoise as the *metabolic network*.

### Evolutionary age of drought-responsive transcriptome in *S. chilense*

To generate a comprehensive understanding of the emergence of the identified drought-responsive GCNs, we estimate the transcriptome ages of the identified cell cycle and fundamental metabolism networks. For that, we build phylostratigraphic profiles for all genes of the two GCNs, summarizing the gene emergence in 18 stages of plant evolution or phylostrata (PS): PS1 representing the emergence of oldest genes (at the time of the first cellular organisms) to PS18 for the most recent genes (*i.e.* present only in *S. chilense*). The PS18 shares no homologue genes with any other species in the nr (non-redundant protein) databases of NCBI (Figure 4A and 4B, Dataset S6). Most genes in the two analyzed GCNs (76.79% in metabolic network and 65.45% in cell-cycle network) are assigned to three main PS: Cellular organisms (PS1), Land plants (Embryophyta; PS5) and Flowering plants (Magnoliopsida; PS8) (Figure 4A). This suggests that the two drought-responsive GCNs we identify have an ancient origin and the components are fairly conserved across the tree of life/plants. Therefore, many drought-responsive pathways likely emerged during the colonization of land by plants (PS5), but many others could derive from exaptation processes from GCNs involved in the core cell process (PS1) or reproductive organ differentiation of flowering plants (PS8). Interestingly, the cell-cycle network shows older origin ages (with more genes (43.73%) assigned to the PS1-3), while the metabolic network presents a larger proportion (48.52%) of genes originating in PS8 (Figure 4A and 4B). Under drought conditions, we also find that cell-cycle network genes of almost all PS ages are down-regulated, while genes of the metabolic network are up-regulated (Figure S5).

Furthermore, we estimate the age of cell-cycle and metabolic GCNs using the transcriptome age index (TAI). We do not find a significant difference of TAI between control and drought samples based on 1,000 randomly selected genes from non-drought responsive genes (Figure S6A; Kolmogorov-Smirnov test, *P* = 0.34), while in cell-cycle and metabolic networks, the mean evolutionary ages of the transcriptomes are significantly different between drought and control conditions (Figure 4C; Kolmogorov-Smirnov test, *P* = 0.03). The TAI profile would be expected to be a flat horizontal line if genes’ ages remain constant across the transcriptomes. In addition, a higher TAI value implies that evolutionary younger genes are preferentially expressed at the corresponding condition/developmental stage. We observe higher TAI in drought samples, supporting that the drought-responsive genes exhibit a younger transcriptome age than genes expressed under control conditions. Moreover, TAI of the metabolic GCN is significantly higher than the cell-cycle (Figure 4C; Kolmogorov-Smirnov test, *P* = 12.51e-7), supporting the previous result that transcriptome ages of the genes in the cell-cycle are older than in the metabolic GCNs.

The contributions of the different PS to the TAI profiles also show notable patterns between the cell-cycle and metabolic GCNs (Figure 4D and 4E). On one hand, early divergent genes (PS1 to PS7) show more constant transcriptome age in all conditions and the genes with ages in PS1, PS5 and PS8 appeared as remarkably important in two GCNs. On the other hand, late-emerging genes (PS8 to PS18) contribute increasingly with their age to the differential expression patterns between control and drought samples, indicating that younger drought-responsive genes are differentially expressed under drought stress in both GCNs (as observed in Domazet-Lošo and Tautz 2010; Piasecka et al. 2013). Remarkably, the youngest genes in PS18 (only found in *S. chilense*), also present a higher contribution in the metabolic GCN, suggesting that these genes are involved in either speciation or local adaptation of *S. chilense* to drought conditions. Note that younger genes (PS9 to PS18) in the cell-cycle GCN hardly contribute to the TAI profile (Figure 4D and 4E).

### Divergence of the drought tolerance transcriptome in *S. chilense*

To drill down into the evaluation of the drought-response mechanisms at the species level, we calculate the TDI index, which represents the mean sequence divergence of a transcriptome. A total of 10 divergence strata (DS) are constructed based on the sequence divergence between genes of *S. chilense* and *S. pennellii* by computing the Ka/Ks ratio (Figure 5A; Figure S7; Dataset S6). The distributions of the Ka/Ks ratio per gene for both GCNs indicate the action of purifying selection, which confirms the conservation of most of drought-responsive genes at the species level. Consistent with the phylostratigraphic patterns, the purifying selection signals in the cell-cycle GCN (Ka/Ks = 0.279 ± 0.333) are higher than in the metabolic GCN (Ka/Ks = 0.329 ± 0.331) (Kolmogorov-Smirnov test, *P* = 2.34e-11; Figure 5A; Table S5). In addition, higher TDI values are observed in the drought samples (Figure 5B) suggesting that the expressed genes we identify in the two GCNs exhibit a more conserved transcriptome profile under control condition compared to drought condition (Kolmogorov-Smirnov test, *P* = 0.004). No significant difference is found between control and drought samples based on 1,000 random genes (Kolmogorov-Smirnov test, *P* = 0.17; Figure S6B). This result supports that different selective pressures act on *S. chilense* GCNs across conditions. In accordance with the TAI results, the transcriptome of the metabolic GCN appears to exhibit a higher transcriptome divergence than the cell-cycle GCN (Figure 5B; Kolmogorov-Smirnov test, *P* = 2.25e-7). Moreover, the low TDI in the cell-cycle GCN and larger TDI differences between drought and control transcriptomes also suggest that regulation of the cell-cycle is likely an ancestral (older) strategy of stress response, not involved in the speciation process. The transcriptome of the cell-cycle GCN may have been evolving and changing in older times, and reached a conserved structure in recent times. Conversely, changes of metabolic pathways and rewiring of the metabolic GCN may appear to be more pronounced and/or common in recent times.

The contributions of the low divergence DS classes (low Ka/Ks in DS1 to DS5) in the cell-cycle GCN (∼ 50% of the genes) are larger than in the metabolic GCN (DS1 to DS5 about 30%), especially in DS1 (lowest Ka/Ks ratio; Figure 5C and 5D). This indicates that purifying selection is acting on genes of the cell-cycle GCN, possibly constraining further changes. In contrast, the metabolic network genes show about 70% contributions in high DS (higher Ka/Ks ratio in DS6 to DS10), especially in DS10 (highest Ka/Ks ratio), indicating that genes in the metabolic network evolve under weaker purifying selection and that recent evolutionary changes occurred.

As a summary, from the deep phylogenetic to the species level, the TAI profile of the cell-cycle network is mainly composed of older phylostrata (PS1 to PS8), while new genes contribute about 20% to the TAI profile of the metabolic network (Figure 4D and 4E). This indicates that the gene expression levels of the cell-cycle network have likely been optimized and fixed early on during evolution, while being maybe also involved in other functional pathways than drought response (Harrison et al. 2012). TDI profiles support this claim: conserved genes do contribute more to the TDI profiles in cell-cycle networks and show adaptive changes in expression for drought response (higher TDI difference between control and drought transcriptomes in cell-cycle network, Figure 5B). In contrast, drought-responsive genes in metabolism network appear more variable in their expression in response to drought stress, because this strategy may be linked to an initial response to severe water scarcity (Dubois and Inzé 2020).

### Population genetics analysis of drought-responsive networks

We also study the selective forces acting on the identified drought-responsive gene networks at the population level. Using full genome sequences of six *S. chilense* populations (C_LA1963, C_LA3111, C_LA2931, SC_LA2932, SC_LA4107, and SH_LA4330; five plants each) recently reported in Wei et al. (2023), aligned to the reference genome of *S. chilense*, we identify 45,208,263 high-quality single-nucleotide variants (SNPs), in which 111,606 SNPs are found in genes of the cell-cycle GCN and 167,334 SNPs in genes of the metabolic GCN. We first compare population structure between the whole-genome data and drought-responsive genes (Figure S8). The results corroborate the genetic structure revealed in Wei et al. (2023) (Figure S8A and S8C). However, the structure exhibited by drought genes shows stronger differentiation among populations than the WGS data (especially for clustering of populations of the central region and SH_LA4330). Moreover, the strong differences from WGS data between the two south coastal populations (SC_LA2932 and SC_LA4107) is attenuated when analyzing SNPs from the drought-responsive genes (Figure S8B and S8D).

We find that the mean nucleotide diversity (π) per gene does not differ between the two GCNs (Figure S9A; Table S5; Kolmogorov-Smirnov test, *P* = 0.15). In addition, the π values of the promoter regions (here 2kb upstream of the transcription initiation site) are significantly higher than those of the gene (coding) regions (Figure S9A; Table S5; Kolmogorov-Smirnov test, *P* = 0.03). This result suggests that the selective constraints in promoter regions may be more relaxed, which could in part explain why certain transcription factors are able to bind to multiple genes in the GCNs. (Table S2). TFs are indeed conserved at the coding sequence level, especially at the functional domains, but higher amount of polymorphism of TF binding sites in the promoter can be indicative of complex and diverse regulation, for example in response to stressful conditions (Spivakov 2014; Sato et al. 2016). Albeit, there is no difference in the nucleotide diversity at the promoter regions between the two GCNs (Figure S9A; Table S5). Furthermore, the genes for the metabolic GCN show lower Tajima’s D values than those of the cell-cycle GCN (Figure S9B; Table S5; Kolmogorov-Smirnov test, *P* = 0.04), suggesting recent positive selection pressure in the metabolic GCN. We find weak correlation between Tajima’s D and Ka/Ks ratio for the cell-cycle GCN and absence of correlation for the metabolic GCN (Figure S10A and S10B). As a negative correlation between Tajima’s D and Ka/Ks ratio is indicative of recent positive selection, our results suggest the possibility of recent positive selection acting on multiple genes within the metabolic GCN (Figure S9B; Table S5).

We further find significant, but opposite, correlations between π or Tajima’s D and the contributions of the different DS for the two GCNs (Figure S10A and S10B). In the cell-cycle GCN, the contributions of different DS have significant positive correlation with π and Tajima’s D (Figure S10A and S10C). This indicates that DS of high contribution to TDI profiles show high nucleotide diversity (and positive Tajima’s D), meaning that older genes are under stronger purifying selection than younger genes in this network because the sequence divergence of cell-cycle genes occurred at old time periods. In contrast, a negative correlation is observed between the contribution of each DS and π or Tajima’s D in the metabolic network (Figure S10B and S10D). Hence, DS with high contribution show low nucleotide diversity and low Tajima’s D, especially DS10. Therefore, it appears likely that the metabolic genes, likely recently evolved, may be under positive selection underpinning the recent evolution of the drought response transcriptome.

### Drought-responsive genes under positive selection promote adaptive evolution in response to drought stress

Genetic drift or changes in selective pressure is one of the main factors that contribute to gene-expression variation (Koenig et al. 2013). To investigate drought-responsive genes that have potentially undergone a shift in selection regime, we search for overlap between genes of two drought-response GCNs studied here and our previously identified 799 candidate genes under positive selection in six populations of *S. chilense* (Wei et al. 2023). We find 74 and 126 drought-responsive genes in the cell-cycle and metabolic networks, respectively in the list of candidate genes under positive selection (Figure 6A; Table S6). These genes exhibit the typical characteristics of positively selected genes with low π and Tajima’s D (Table S5). This indicates that drought stress is likely an important driver of adaptation and these drought-response genes may play key roles for colonization of new arid habitats. Similar numbers of drought-responsive genes likely under positive selection are observed across different populations of *S. chilense* encompassing different parts of the range, except for SH_LA4330 (Wei et al. 2023). The number of candidate genes belonging to the metabolic or cell-cycle GCNs is similar in the three central populations (C_LA1963, C_LA3111 and C_LA2931) (Figure 6A; Table S6). The most recent diverged highland population (SH_LA4330) contains the largest number of positively selected drought-responsive genes (Figure 6A; Table S6) with a similar proportion of genes from both networks. Noticeably, in the two south-coast populations (SC_LA2932 and SC_LA4107) a large majority of genes under positive selection belong to the metabolic GCN (showing absence of cell-cycle genes in population SC_LA2932, Figure 6A; Table S6).

Previous studies have demonstrated that adaptive genes are pleiotropic and proposed functional connectivity between networks related to different quantitative traits (Wagner et al. 2007; Erwin and Davidson 2009; Hämälä et al. 2020). To address the role that (putatively) positively selected genes may play within the drought-responsive networks, we compare the connectivity of these genes in the two networks (Figure 6B; Table S7). In the metabolic network, the connectivity of positively selected genes (0.55 ± 0.10) is significantly higher than other drought-responsive genes (0.44 ± 0.12) (Figure S11A; Kolmogorov-Smirnov test, *P* = 0.017), but we do not observe such significant difference for the cell-cycle network (Figure S11A; Kolmogorov-Smirnov test, *P* = 0.43). Furthermore, the connectivity of positively selected genes of the metabolic network is much higher than those from the cell-cycle network in six populations (Figure 6B; Table S7; Kolmogorov-Smirnov test, *P* = 0.007). These results suggest that highly connected (likely more pleiotropic) genes in the metabolic GCN may have facilitated the recent colonization of new habitats (Hämälä et al. 2020) during the divergence process of *S. chilense*. In contrast, the connectivity of positively selected genes in the cell-cycle network is significantly lower (Figure S11A). Therefore, we suggest that the two networks underwent different evolutionary selective pressures during the range expansion of *S. chilense*.

Finally, we compare the age of the selective sweep at the candidate genes of the two GCNs based on the results in Wei et al. (2022). We find that sweep ages at the cell-cycle genes are slightly younger than at those of the metabolic network, especially in the three highland populations (C_LA2931, C_LA3111 and SH_LA4330; Figure S11B and S11C; Table S7). This supports that drought adaptation is an important mechanism underlying the recent (re)colonization of highland habitats (Raduski and Igić 2021; Wei et al. 2023). Interestingly, we find significantly positive correlation between the age of the sweep and gene connectivity for both GCNs and across all six populations (Figure 6C). Figure 6D and 6E provide the visualizations of two networks and exhibit the relationship between sweep age and connectivity (depicting weighted connection strength greater than 0.65 between any two genes). In other words, it appears that selective sweeps occur first at more connected genes and, subsequently at less connected genes, during the history of colonization/adaptation of new arid habitats. To our knowledge, this is the first report of a correlation between the age of a selective sweep and the connectivity of genes in a network. To obtain more evidence to support this inference, we also calculate the tMRCA (time to most recent common ancestor) to estimate the age of drought-responsive genes based on allele frequency of SNPs. The positive correlation between tMRCA of drought-responsive genes under the positive selection and connectivity is also supported (Pearson’s cor=0.69, *P* = 2.47e-5), consistent with the correlation with sweep age. Moreover, the low correlation (Pearson’s cor=0.31, *P* = 0.14) is observed between tMRCA of other (outside of sweep regions) drought-responsive genes and connectivity. This supports the hypothesis of polygenic adaptation in GCNs where the positive selection acts first on core genes (with high connectivity and more pleiotropic) of networks, and subsequently on the peripheral genes (less connectivity and less pleiotropic). These positively selected genes ultimately regulate the expression of other genes in the network.

## Discussion

In this study, we identify two drought-responsive GCNs by analyzing gene expression profiles of plants growing under control and drought conditions. Two GCNs involved in cell-cycle and metabolic biological processes are detected and their structural relevance are supported by TF/TFBS predictions. These networks represent two different strategies for drought response (Farooq et al. 2009; Danilevskaya et al. 2019). We then demonstrate that the cell-cycle network is evolutionary older and more conserved than the metabolic network. Despite the ancient history of these two GCNs, we further show that both GCNs also contribute to different extents to contemporary processes of adaptation to drought conditions when *S. chilense* colonizes new arid habitats around the Atacama desert. The joint analyses of genomic and transcriptomic data indicates that 1) at the transcriptome level, metabolic GCN present a higher evolvability especially with younger selection events linked to response to new environments, 2) cell-cycle GCN is less evolvable, and 3) both networks still present signals of evolution under positive selection in core elements of the GCN, while peripheral genes of the network can be involved in adaptation at later stages of the colonization processes.

### Drought tolerance is mediated by regulation in cell proliferation and metabolism

When roughly defining the organ development into cell proliferation and differentiation, water deficit appears to be a limiting factor for both processes (Alves and Setter 2004; Verelst et al. 2013). Drought stress reduces the activity of the cell cycle and thus slows down the growth and development of plants. The down-regulated genes we find in the cell-cycle network also indicate that genes related to cell cycle are suppressed by drought stress possibly to restrict the cell division in *S.chilense.* Reduction of cell number due to mild drought stress is also found in *A. thaliana* (Skirycz and Inzé 2010). This means that the cell-cycle response to drought may be very general and indirect. However, our speculations are mainly based on the aboveground tissues of *S. chilense*. Conversely, the changes of fundamental metabolic activity may be a faster and a flexible drought-responsive strategy presumably related to acclimation (Harb et al. 2010). Plant water shortage is first reflected in changes in metabolic processes, such as accelerating the catabolism of macromolecules in order to regulate the penetration of tissues, to maintain physiological water balance, or slowing down metabolism to reduce energy and water consumption (Reddy et al. 2004; Gupta et al. 2020). In addition, the signaling pathways related to the metabolic gene network are also demonstrated to be a response to drought stress, for example, the abscisic acid (ABA) signaling pathway regulates the response to dehydration and optimizes water utilization (Harb et al. 2010; Wilkinson and Davies 2010). Although these two GCNs correspond to two different strategies of drought response, they are not isolated, but interact with one another in a time-dependent manner. Water deprivation and heat first change the metabolic processes leading to stomata closure, which leads then to cell cycle network to be affected under long-term lack of water. In return, the increased or decreased cell cycle gene expression affects the further physiology and metabolism of the plant (Gupta et al. 2020). Indeed, drought-responsive strategies regulating the cell cycle appear to be activated later than metabolism processes, as glucose metabolism rapidly follows drought stress, whereas the accumulation of amino acids which is a crucial part of the cell cycle response starts at a later time in response to drought (Fàbregas and Fernie 2019).

### Rewiring of ancient GCNs drives recent adaptation to dry environments

The phylostratigraphic analysis supports that the majority of drought-responsive genes in *S. chilense* evolved during the early to middle stages of plant evolutionary history, which is in agreement with the time of origin of multiple abiotic response genes in *Arabidopsis thaliana* (Mustafin et al. 2019). This reinforces that the emergence of drought-responsive genes coincides with the time periods of divergence among major plant groups (land- and flowering plants), which are marked by frequent whole genome duplication events that trigger gene family expansions, gene neo- and sub-functionalization, and genome reorganization processes (Wang et al. 2012; Clark and Donoghue 2018). These genomic processes likely contributed to the enrichment of drought-responsive GCNs. For instance, fundamental morphological traits involved in drought responses, such as stomata, are present in the ancestral land plants. However, stomatal genes existed prior to the divergence of land plants and underwent multiple duplications during the course of evolution. Additionally, their response to environmental cues, such as humidity, light, CO2, and ABA, is widely distributed and may be ancestral to land plants (Clark et al. 2022). Therefore, we propose that our two drought-responsive networks were primarily established during or shortly after the divergence of land plants and have subsequently undergone expansion. This highlights the crucial role of ancestral genomic processes in shaping the genetic mechanisms that underlie plant adaptation to drought.

Previous studies show that TAI and TDI profiles across embryogenesis, seed germination and transition to flowering in *A. thaliana* exhibit a ‘hourglass pattern’ (older and conserved transcriptomes are preferentially active at the mid-development stages; Quint et al. 2012; Drost et al. 2016). However, our TAI/TDI profiles for the two developmental stages remain stable under the same conditions (Figures 4C and 5B). The similar TAI/TDI between developmental stages (Figure 4C and 5B) that we obtained is certainly because our analyses focused on two modules (co-expressed genes) highly correlated to the differential expression between drought and control conditions (Figure 2D; Table S1). Therefore, developmental stage-specific response genes are underrepresented in the two analyzed networks. However, increased TAI/TDI values under drought conditions suggest that stress response transcriptomes are composed of relatively more recently diverged genes, and therefore are more evolvable. We suggest that this inference needs to be verified in other stress responsive transcriptomes (salt, heat, cold, etc.). We then speculate, that although abiotic stress response regulatory networks are mostly composed of highly ancient and conserved elements across species (Chen and Zhu 2004), networks retain the ability to change expression patterns to respond rapidly to environmental changes or to explore new ecological niches. Moreover, given the pleiotropic nature of the abiotic stress-response traits, we can expect shared patterns of evolution (at the constitutive and expression components) of the networks for different stress conditions (and possible trade-offs between traits and GCNs).

Extensive network rewiring in relatively recent and short time-frames have been found in maize and tomato in response to domestication (Swanson-Wagner et al. 2012; Koenig et al. 2013). It is therefore not surprising to find signs of adaptive variation in core elements of rather conserved regulatory networks related to the colonization processes of new (here arid) habitats. The genetic (and morphological) divergence of the *S. chilense* marginal southern populations, southern coastal and highland, is recent but strong (Raduski and Igić 2021). It is congruent with theoretical results showing that gene networks with higher mutation sensitivity can facilitate local adaptation, present increasing variance in gene expression and underlie accelerated range expansion processes across abiotic environmental gradients (Deshpande and Fronhofer 2022). Complementarily, our empirical approach shows the existence of two regulatory networks with different evolutionary trends, one being more conserved than the other and exhibiting different gene expression responses. One GCN would exhibit a faster and more variable response (metabolic), while the other a later (delayed) but more constitutive response (cell-cycle) to drought. Despite the differences in gene age and variation between the networks, our results show that both GCNs have undergone sufficient changes leading to their rewiring during the divergent process of colonization of *S. chilense* around the Atacama. Nevertheless, genes in the metabolic network show more recent evolution, with new genes members appearing in *S. chilense*, concomitantly with more variable expression in the drought transcriptome.

These drought-responsive genes to *S. chilense* likely facilitated the adaptation of this species to unique arid (up to hyper-arid) habitats, especially when colonizing the southern part of the range. Indeed, population structure based on SNPs indicates that drought-responsive genes reflected adaptation/colonization to arid habitats in *S. chilense* (Figure S8). Importantly, we found about 200 drought-responsive genes previously identified as candidate genes under positive selection (*i.e.* located within sweep regions in Wei et al. 2023). This confirms that drought stress is an important driver of ecological divergence in *S. chilense.* We finally provide some indirect evidence that changes at central genes (with higher connectivity) can be responsible for the short-term response to selection (Jovelin and Phillips 2009; Luisi et al. 2015) and promote rewiring of the gene network (Koubkova-Yu et al. 2018). Thus, highly connected genes may be targets of positive selection during the first phase of the environmental change or colonization to contrasting environments, and may be keys for ‘piggybacking’, defined as the change in gene expression of a focal gene driving phenotypic change. Altogether, our results on the age-dependent adaptive role of genes with different network connectivity (and possible pleiotropic effects) provide another line of evidence supporting the view that molecular evolution follows an adaptive walk and are complementary to the recent study by (Moutinho et al. 2022).

### Limitations and further work

A limitation of our gene expression study is that our transcriptomic analyses are based on individuals from a single location (near the putative region of origin of the species; Wei et al. 2022), while variability in gene expression and phenotypic response has been observed between different populations (Mboup et al. 2012; Fischer et al. 2013; Nosenko et al. 2016). Further expression studies including plants from multiple locations would be useful to verify that the identified GCNs are also present and expressed in other populations and study the possible variation in the most southern populations. More evidence based on multiple populations is needed to confirm the ‘piggybacking’ phenomenon of gene expression in *S. chilense*. Additional support on the variability of transcriptome evolution across populations as well as long read sequencing of more genomes will be beneficial in assessing the role of gene duplication and gene deletion yielding the evolution of the gene networks. Such studies would also allow the analysis of evolution of adaptive gene networks and polygenic selection occurring for complex traits such as drought tolerance. Finally, more detailed studies with a larger sample size from the field will help to discover other gene networks and their interactions related to abiotic stress and the evolution of the species. A detailed discussion of the potential biases associated with the use of multiplied accessions at TGRC (Tomato Genetics Resource Center, UC Davis, USA) compared to samples from natural populations is found in Wei et al. (2022). Sampling and experimental work in the field would improve the resolution of transcriptome and genomic studies, in order to assess phenotypic differences between organs and stages of development and thus extend the knowledge to other relevant characteristics such as secondary metabolism, which is known to have relevant influence on biotic and abiotic interactions (Mes et al. 2008; Bolger et al. 2014; Tapia et al. 2022).

## Material and methods

### Plant material and drought stress experiment

Seeds of *S. chilense* accession LA1963 were acquired from Tomato Genetics Resource Center (TGRC), University of California at Davis. Seeds were soaked in 50% household bleach (2.7% sodium hypochlorite) for 30 minutes and rinsed thoroughly with water according to instructions provided by TGRC. The rinsed seeds were sown into pots containing sterilized soil with perlite and sand (1:2) and grown under controlled conditions (22C day/20C night, 16h light/8h dark photoperiod). On the 24th day after sowing, all plants were randomly distributed into two groups and watered with a sufficient volume to reach the bottom of containers (30-40 ml). The first group of plants were maintained under normal watering condition, watered with a sufficient volume of water (50-55 ml) on 4, 7 and 11 days after start of the experiment (day 24). A moderate water stress regime was imposed to second group of plants by stopping irrigation for 7 days followed by re-watering with 25 ml of water. On day 12, newly expanded leaf (1-1.5 cm length) and shoot apices with immediately surrounding leaf primordia (shoot apices and P1-P5 leaf primordia) from each group were dissected carefully using razor blades and immediately grounded into fine powder in liquid nitrogen for RNA extraction. Four biological replicates were used for all RNA-Seq experiments from each tissue type. Each replicate of leaf and shoot apex samples included the pooled tissues from five and six plants, respectively.

### RNA extraction and cDNA library construction

Libraries were constructed and named as follows: leaves under control (optimal watering) condition (CL-A to D), shoot apices under control condition (CSA-E to H), leaves under drought condition (DL-I to L), and shoot apices under drought condition (DSA-M to P). Tissues were lysed using zircon beads in Lysate Binding Buffer containing Sodium Dodecyl Sulfate. mRNA was isolated from 200 μl of lysate per sample with streptavidin coated magnetic beads for indexed non-strand specific RNA-Seq library preparation according to the method described by (Kumar et al. 2012). 1 μl of 12.5 μM of 5-prime biotinylated polyT oligonucleotide and streptavidin-coated magnetic beads were used to capture mRNA and isolate captured mRNAs from the lysate, respectively. Equal amount of mRNA of each experimental group were used to construct 16 libraries. For library construction the rapid version of Kumar et al. (2012) RNA-sequencing method (Townsley et al. 2015) was used. Each sample was barcoded using standard Illumina adaptors 1-16 to allow up to 16 samples to be pooled in one lane of sequencing on Illumina HiSeq4000. The libraries were eluted from the pellet with 10 μl 10 mM Tris pH 8.0 and pooled as described by Kumar et al. (2012). Quantification and quality assessment of resulting libraries were performed on Fragment Analyzer (FGL_DNF-474-2-HS NGS Fragment 1-6000bp.mthds) and sequenced using the Illumina HiSeq 4000 platform to generate 100 bp single-end reads at the Vincent J. Coates Genomic Sequencing Facility at UC Berkeley.

### Transcriptome and genome data processing and mapping

For transcriptome data, the adapters were removed from raw reads by two consecutive rounds using BBDuk in BBTools v38.90 (Bushnell 2014). Two sets of parameters were used in two rounds respectively: first round ‘ktrim=r k=21 mink=11 hdist=2 tpe tbo minlength=21 trimpolya=4’; second round ‘ktrim=r k=19 mink=9 hdist=1 tpe tbo minlength=21 trimpolya=4’. Then Low-quality reads were also removed with BBDuk using parameters ‘k=31 hdist=1 qtrim=lr trimq=10 maq=12 minlength=21 maxns=5 ziplevel=5’. The clean reads of each sample were mapped to the *S. chilense* reference genome (Silva-Arias et al. submited) using BBMap in BBTools. The SAM files were then converted and sorted to BAM files using Samtools v1.11 (Wysoker et al. 2009). The number of reads were mapped to each gene were counted via featureCounts v2.0.1 in each sample (Liao et al. 2014). To eliminate the differences between samples, the gene expression level was normalized using the TPM (Transcripts Per Kilobase Million) method (Wagner et al. 2012).

The relationships among transcriptome samples were evaluated using the TPM values. The correlation coefficient between two samples was calculated to evaluate repeatability between samples using Pearson’s test. Principal component analysis (PCA) was performed using the *prcomp()* function in R (R Core Team 2020) based on TPM values.

### Identification of differentially expressed genes and gene co-expression analysis

Differential expression analysis of groups among the different conditions and tissues was performed using the DESeq2 R package (Love et al. 2014). The raw read counts were inputted to detect Differential Expressed Genes (DEGs). The *P-*value ≤ 0.001, the absolute value of log2FoldChange ≥ 1 and a false discovery rate (FDR) adjusted *P* ≤ 0.001 were classified as differentially expressed genes.

To identify the gene co-expression networks, weighted gene correlation network analysis (WGCNA) was constructed using TPM values to identify specific modules of co-expressed genes associated with drought stress (Langfelder and Horvath 2008). We first checked for genes and samples with too many missing values using *goodSamplesGenes()* function in WGCNA R package. We then removed the offending genes (the last statement returns ‘FALSE’). To construct an approximate scale-free network, a soft thresholding power of five was used to calculate adjacency matrix for a signed co-expression network. Topological overlap matrix (TOM) and dynamic-cut tree algorithm were used to extract network modules. We used a minimum module size of 30 genes for the initial network construction and merged similar modules exhibiting > 75% similarity. To discover modules of significantly drought-related, module eigengenes were used to calculate correlation with samples with different conditions. The visualization of networks were created using Cytoscape v3.8.2 (Su et al. 2014).

### Identification of transcript factor families and transcript factor binding sites

The protein sequences were obtained from the reference genome and annotation ‘gff’ file with GffRead (Pertea and Pertea 2020), and were used to identify TF families using online tool PlantTFDB v5.0 (Guo et al. 2007). Furthermore, the upstream 2000 bp sequences of the transcription start sites (TSS) were extracted as the gene promoter from the reference genome to detect TFBS. The TFBS dataset of relative species *S. pennellii* was also downloaded from Plant Transcriptional Regulatory Map (PlantRegMap, http://plantregmap.gao-lab.org/) as background of TFBS identification (Tian et al. 2020). Then, the TFBS of *S. chilense* was identified using FIMO program in motif-based sequence analysis tools MEME Suit v5.3.2 (Bailey et al. 2015). The TFBS was extracted with p < 1e-5 and q < 0.01.

### Gene ontology (GO) analysis

We first constructed the dataset of assigned GO terms for all genes used protein sequence by PANTHER v16.0 (Mi et al. 2021). Then, the GO enrichment analysis of drought-responsive genes was performed using clusterProfilter v3.14.2 (Yu et al. 2012). Benjamini– Hochberg method was used to calibrate *P* value, and the significant GO terms were selected with *P-*value below to 0.05.

### Construction of phylostratigraphic map

We performed phylostratigraphic analysis based on the following steps. First, the phylostrata (PS) was defined according to the full linkage of *S. chilense* from NCBI taxonomy database. The similar PS was merged and finally 18 PS were generated (Figure 4A). Second, the protein sequences were blast to a database of non-redundant (nr) proteins downloaded from NCBI (https://ftp.ncbi.nlm.nih.gov/blast/db/) with a minimum length of 30 amino acids and an E-value below 10^-6^ using blastp v2.9.0 (Camacho et al. 2009). Third, each gene was assigned to its PS by the following criterion: if no blast hit or only one hit of *S. chilense* with an E-value below 10^-6^ was identified, we assigned the gene to the youngest PS18. When multiple blast hits were identified, we computed lowest common ancestor (LCA) for multiple hits using TaxonKit v0.8.0 (Shen and Ren 2021) and then assigned LCA to specific PS.

### Construction of divergence map

We performed divergence stratigraphy analysis to construct sequence divergence map of *S. chilense* using function *divergence_stratigraphy()* of R package ‘orthologr’ (Drost et al. 2015) following four steps: 1) the coding sequences for each gene of *S. chilense* and *S. pennellii* (NCBI assembly SPENNV200) were extracted from their reference and annotation files. 2) We identified orthologous gene pairs of both species by choosing the best blast hit for each gene using blastp. We only considered a gene pair orthologous when the best hit has an E-value below 10^-6^, the gene pair is considered orthologous; otherwise, it is discarded. 3) Codon alignments of the orthologous gene pairs were performed using PAL2NAL (Suyama et al. 2006). Then, Ka/Ks values of the codon alignments were calculated using Comeron’s method (Comeron 1995). And 4) all genes were sorted according to Ka/Ks values into discrete deciles, which are called divergence stratum (DS).

### Estimation of transcriptome age index and transcriptome divergence index

The TAI is computed based on phylostratigraphy and expression profile, which assign each gene to different phylogenetic ages by identification of homologous sequences in other species (Domazet-Lošo et al. 2007). The evolutionary age of each gene was quantified combining its PS and expression level to obtain weighted evolutionary age. Finally, weighted ages of all genes are averaged to yield TAI, which is defined as the mean evolutionary age of a transcriptome (Domazet-Lošo and Tautz 2010). A lower value of TAI describes an older mean evolutionary age, whereas a higher value of TAI denotes a younger mean evolutionary age and implies that evolutionary younger genes are preferentially expressed in the corresponding sample or condition (Domazet-Lošo and Tautz 2010; Piasecka et al. 2013). The TDI represents the mean sequence divergence of a transcriptome quantified by divergence strata (DS) and gene expression profile (Quint et al. 2012). The genes are assigned to different DS and then weighted by their expression level to yield the TDI. A lower value of TDI describes a more conserved transcriptome (in terms of sequence dissimilarity), whereas a higher value of TDI denotes a more variable transcriptome. Here, we calculate TAI and TDI profiles in different samples using *PlotSignature()* function of the myTAI R package.

### Population genetics analysis and detection of positive selection on drought-responsive genes

Whole-genome sequence data from six populations *S. chilense* (five individuals each) previously analyzed in (Wei et al. 2022; BioProject PRJEB47577) were used to calculate population genetics statistics for coding and promoter region sequences for all genes identified in the GCNs. Single nucleotide variants (SNPs) based on the short-read alignment to the new reference genome for *S. chilense* (Silva-Arias et al. submitted) using the same methods in Wei et al. (2022). Population genetics statistics namely, nucleotide diversity (π) and Tajima’s D were calculated with ANGSD v0.937 (Korneliussen et al. 2014) over gene and promoter regions. These statistics first were calculated at per site in gene and promoter regions, and then we used a R script (https://gitlab.lrz.de/population_genetics/s.chilense-drought-transcriptome) to obtain statistics in each gene and promoter regions. PCA on SNP data from 30 whole genomes was also performed using GCTA (v1.91.4; Yang et al. 2011). The genetic structure inference was performed using ADMIXTURE v1.3.0 (Alexander et al. 2009).

Drought-responsive genes under positive selection were extracted by blast (e-value < 1e-6) between drought-responsive genes identified in this study and the genes located inside sweep regions in our previous study using *S. pennellii* as the reference genome. We also use the sweep ages obtained in Wei et al. (2022).

### Estimation of allele age

We implemented in GEVA (Genealogical Estimation of Variant Age; Albers and McVean 2020) to dating genomic variants in the drought-responsive genes. We generated input for GEVA based on the recombination rate 3.24 x 10^-9^ per site per generation (based on the overall recombination density in *S. lycopersicum* [1.41 cM/Mb] Anderson and Stack 2002; Nieri et al. 2017; and within the possible range of rates used in Wei et al. 2022). We used population size (*N*_e_) 20,000 and mutation rate 5.1 x 10^-9^ (Roselius et al. 2005; Wei et al. 2023), and then relied on the recombination clock to estimate the age of alleles (tMRCA).

## Supplementary material

Supplementary data are available at Molecular Biology and Evolution online.

## Acknowledgements

KW was funded by the Chinese Scholarship Council. SS was funded by the Fulbright Visiting Scholar Program of the U.S. Department of State and by German Academic Exchange Service (DAAD) within the framework of Research Stays for University Academics and Scientists. GAS-A was funded by the Technical University of Munich. AT acknowledges funding from DFG (Deutscheforschungsgemeinschaft) Grant Number: 317616126 (TE809/7-1). NS acknowledges funding from U.S. National Science Foundation (NSF/ IOS-1238243). This work was supported by grants from JSPS KAKENHI (JP19K23742, JP20K06682, JP20KK0340 to H.N.), and NSF PGRP IOS-1856749 and IOS-211980 to NS. The laboratory work of H.N. was supported by a Grant-in-Aid for Scientific Research on Innovative Areas (JP19H05672). We thank the Tomato Genetics Resource Center (TGRC) of the University of California, Davis for generously providing us with the seeds of the accession included in this study.

## Data Availability

The raw sequencing RNA data is available in PRJDB15063 (*S. chilense*), PRJEB5809 (*S. pennellii*) and PRJNA812356 (*S. lycopersicum*). The raw pair-end whole-genome sequencing data can be accessed at the European Nucleotide Archive (ENA) project accession PRJEB47577. All codes used in this study and other previously published genomic data are available at the sources referenced. The code for implementing the analyses used in this paper can be found on our GitLab repository: https://gitlab.lrz.de/population_genetics/s.chilense-drought-transcriptome

---

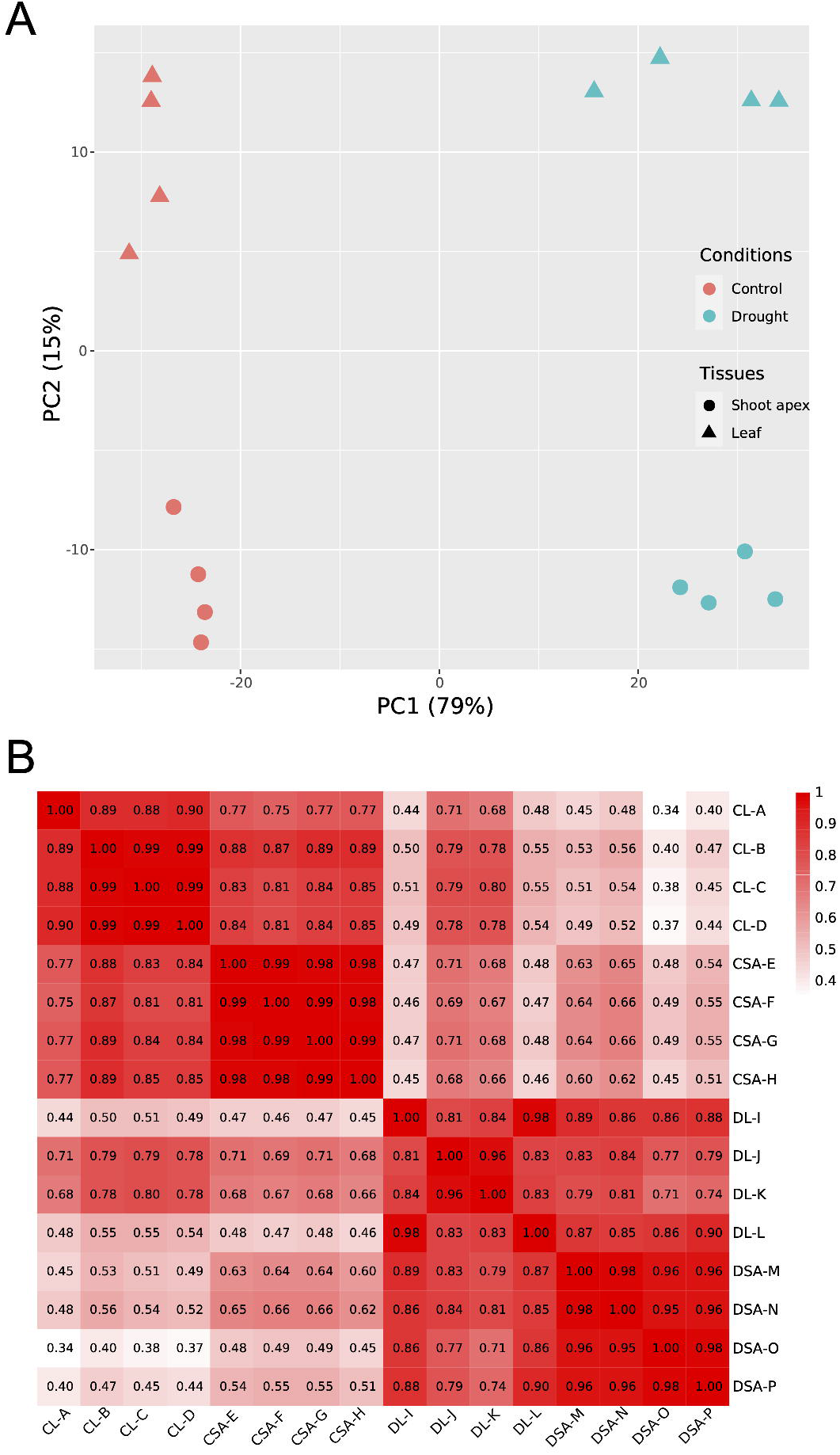

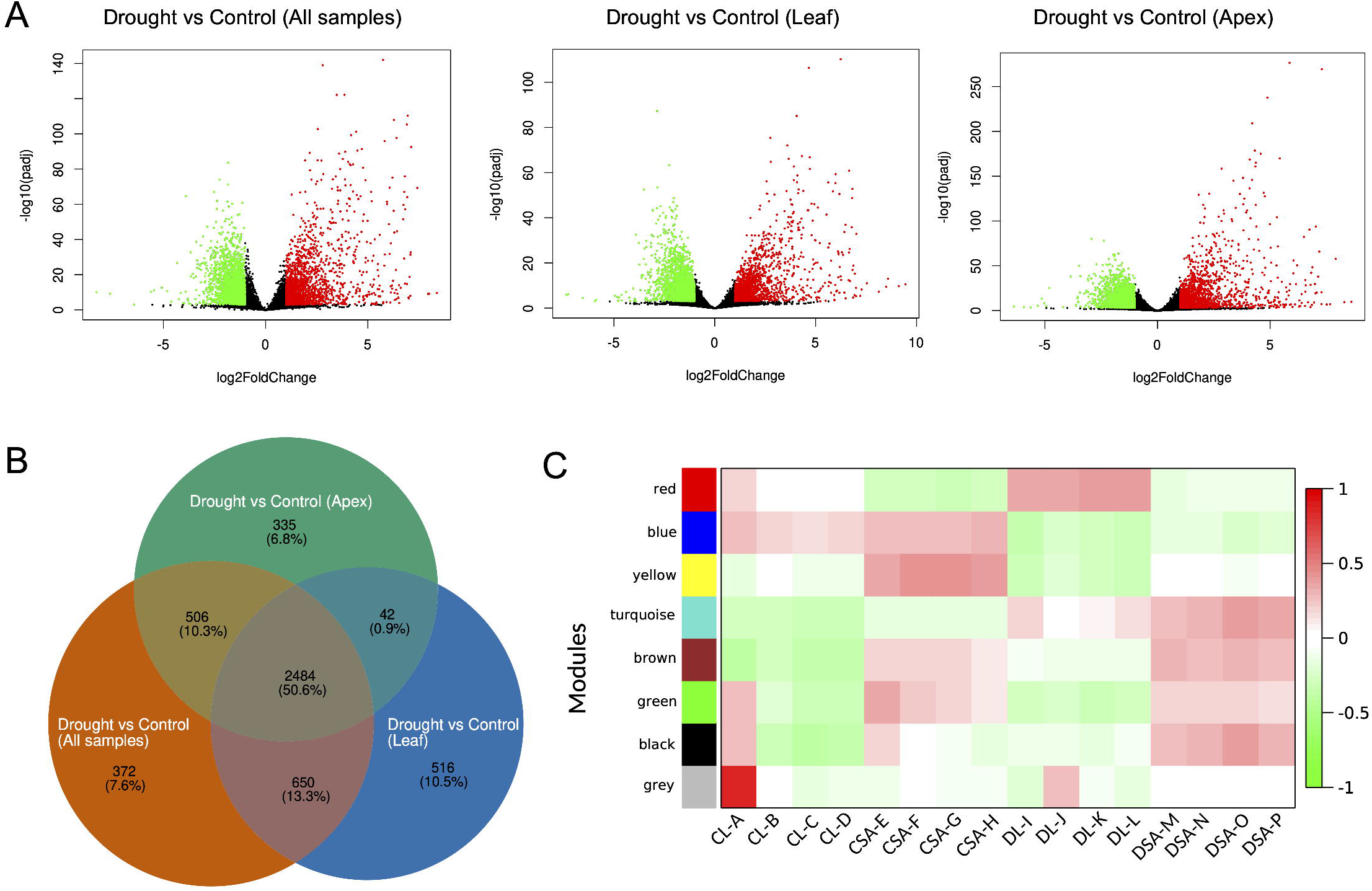

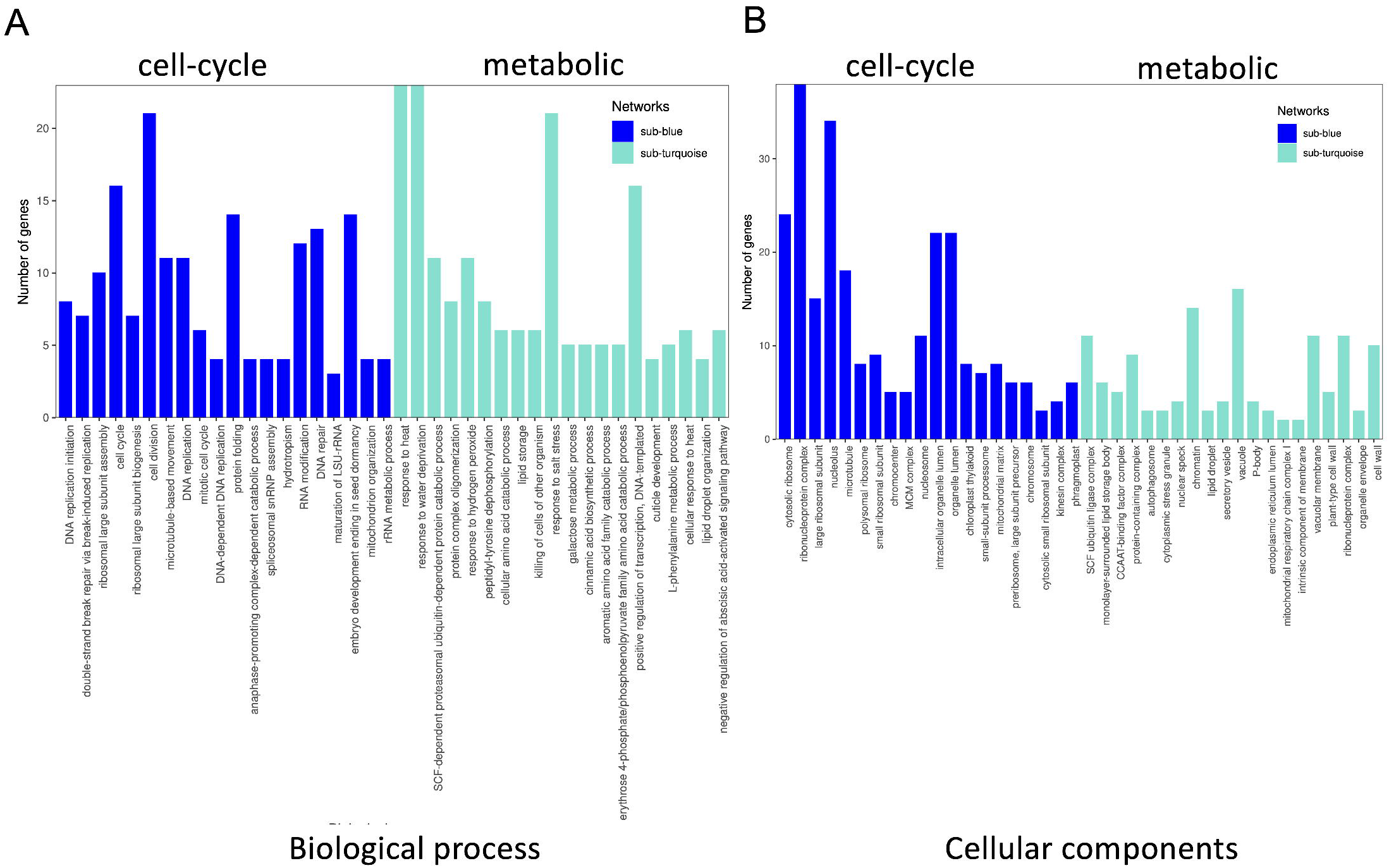

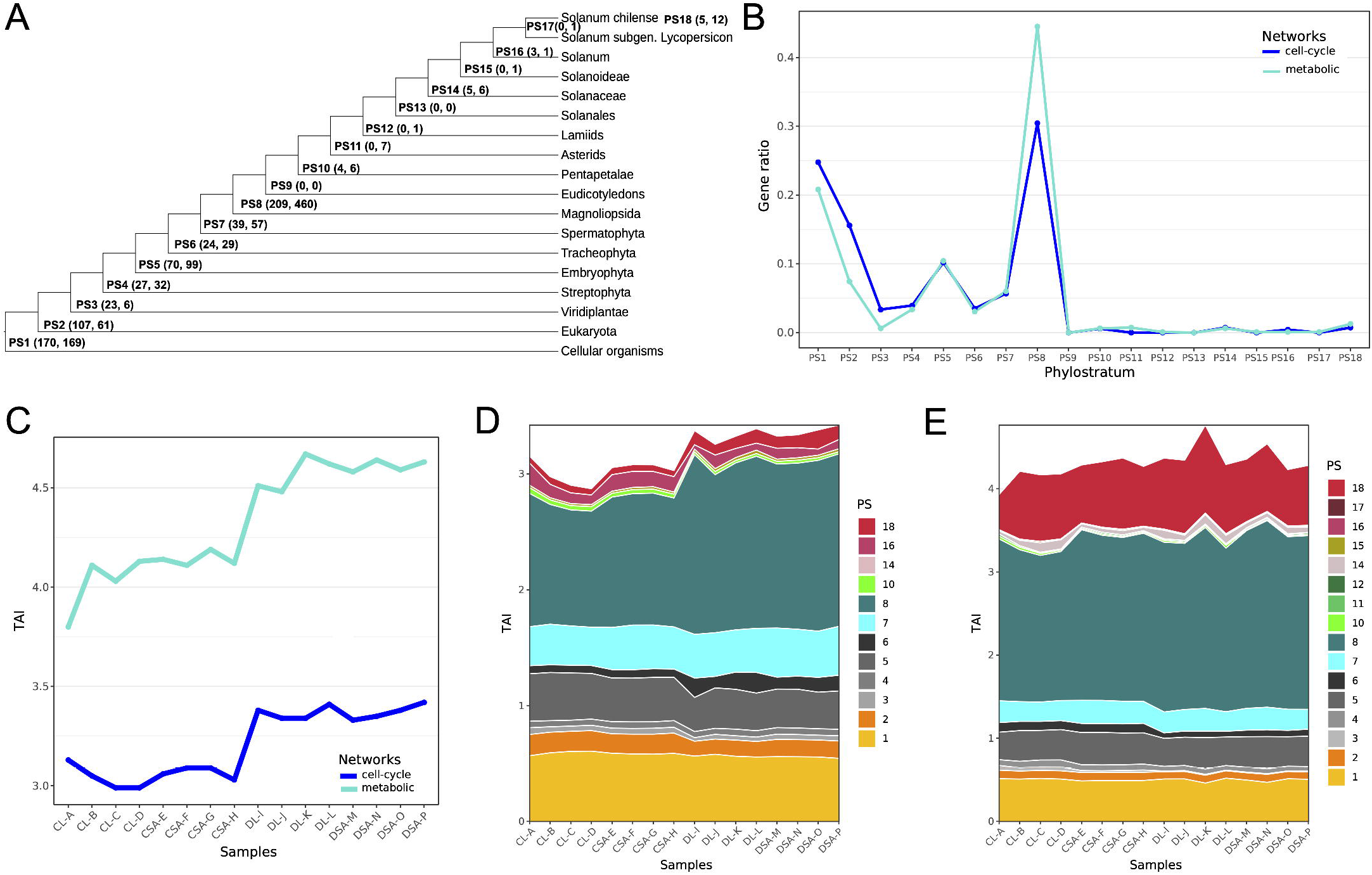

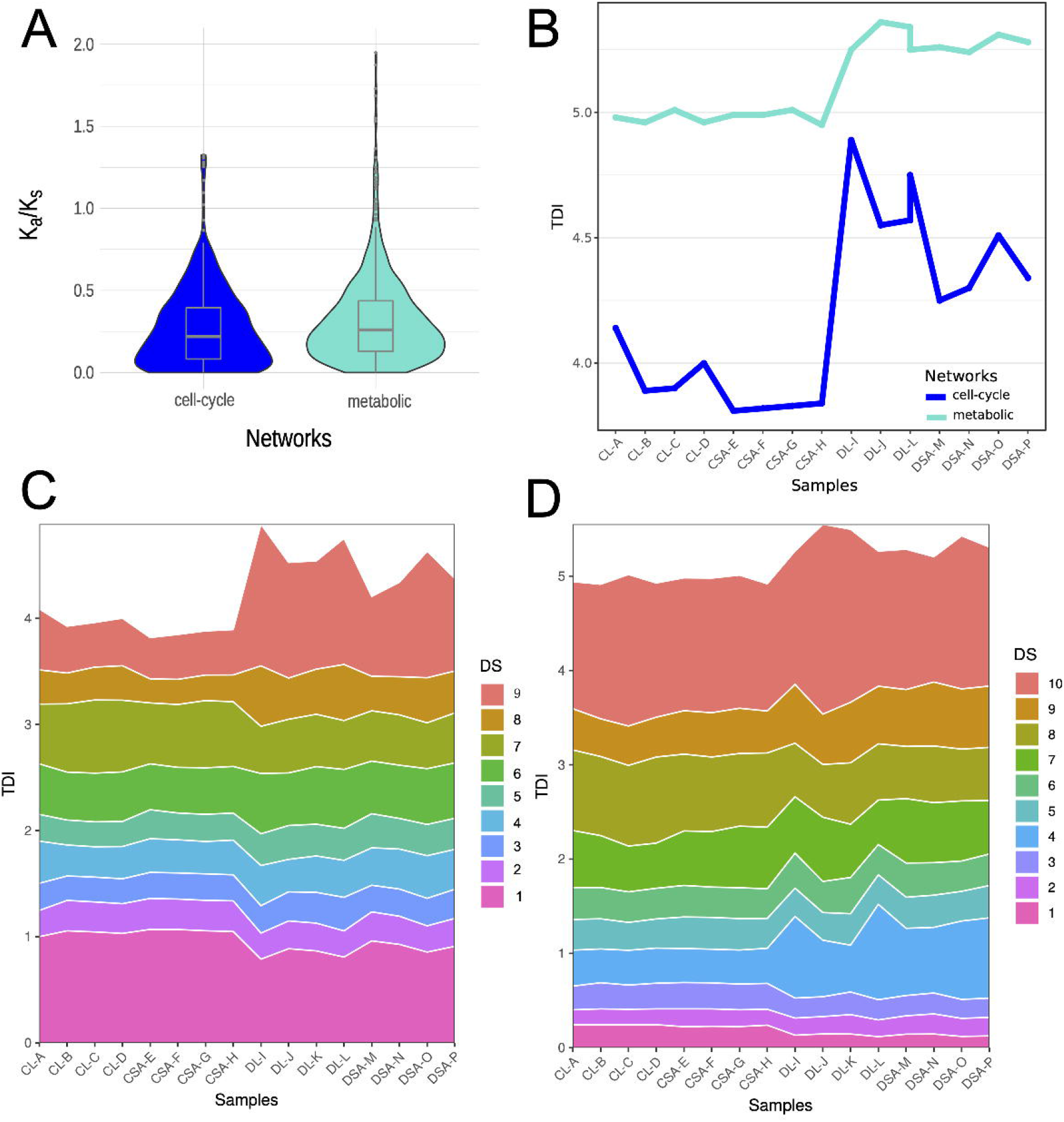

