## Supplementary data for "Evolution of two gene networks underlying adaptation to drought stress in the wild tomato *Solanum chilense*"

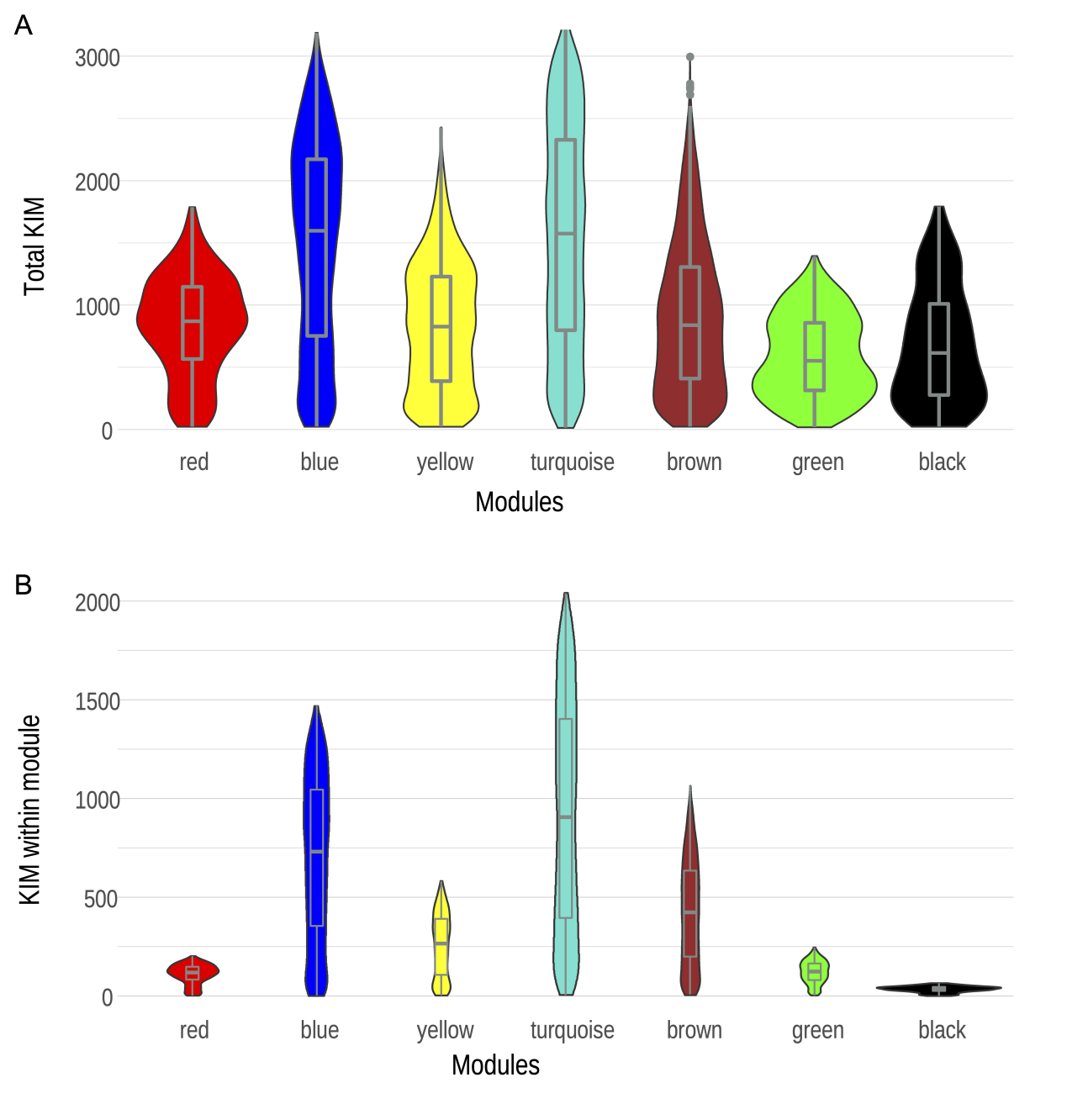

**Figure S1**. Intramodular connectivity measures (KIM) show connection or co-expression of a given gene with respect to the genes. (A) KIM of a given gene with all genes used to WGCNA. (B) KIM of a given gene with genes in the specific network. We focus thereafter on the blue (defined as cell-cycle) and turquoise (defined as metabolic) modules.

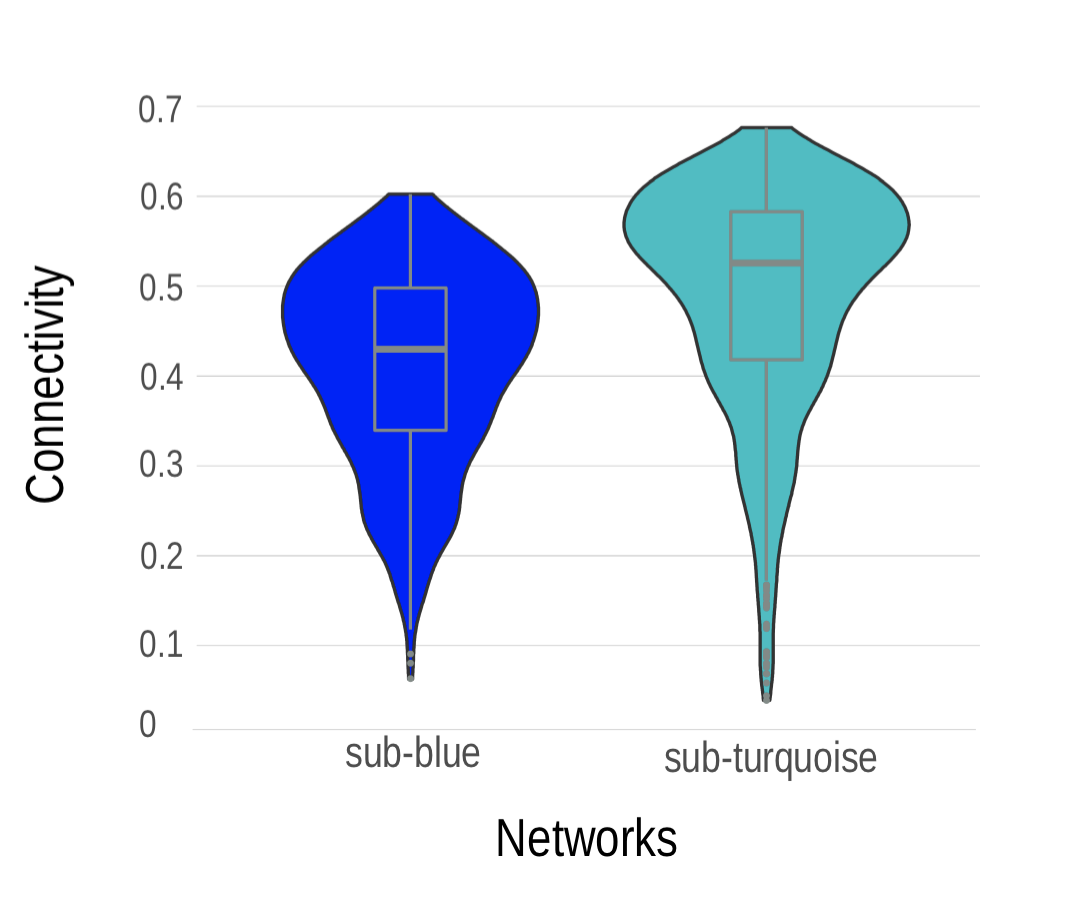

**Figure S2**. Relative connectivity of the reconstructed sub-blue (cell-cycle) and sub-turquoise (metabolic) network. The values were calculated as KIM divided by number of genes within each specific module.

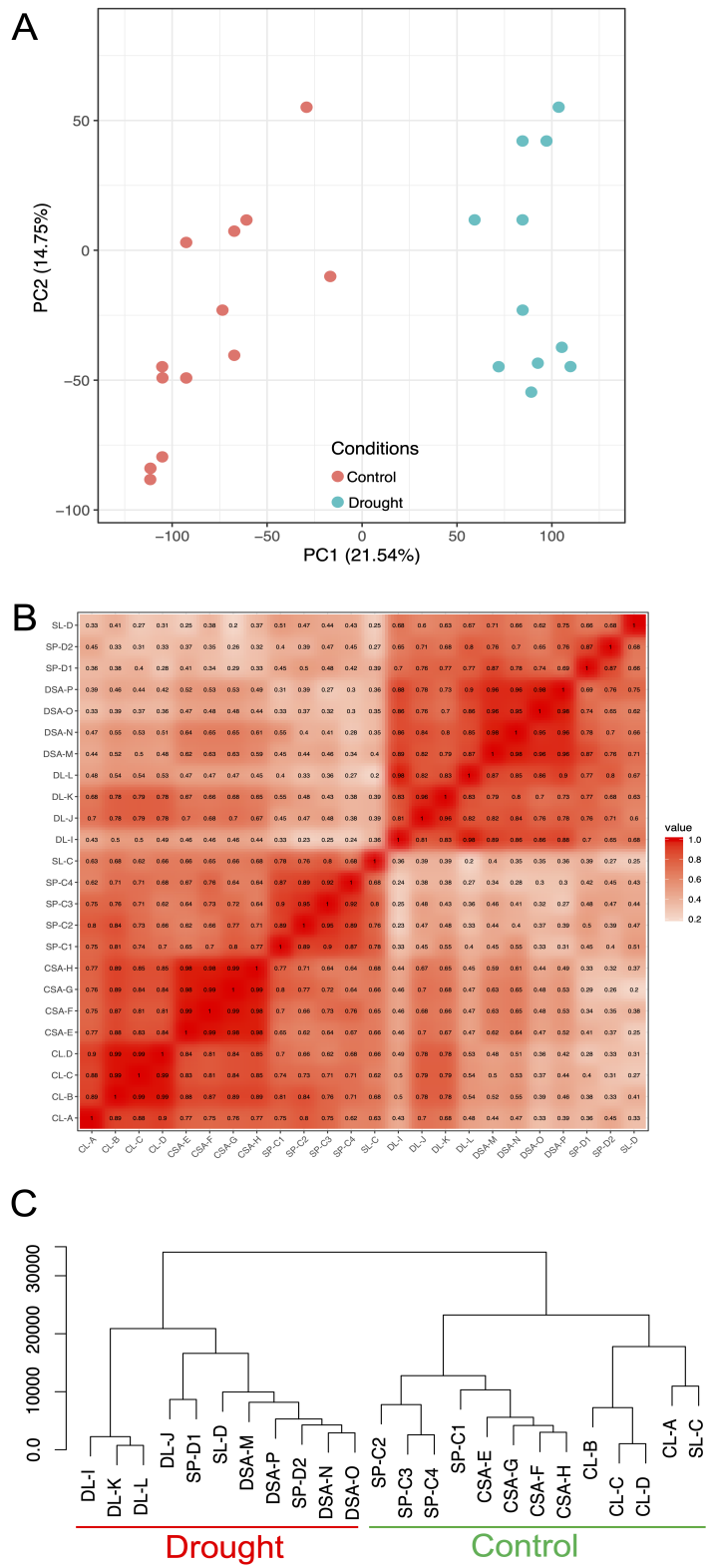

**Figure S3.** Exploratory analyses of RNA-seq differential expression patterns in 24 libraries of *S. chilense*, *S. pennellii* and *S. lycopersicum*. (A) PCA reveals stronger clustering associated with the experimental conditions. (B) Heatmap plot of sample correlation (Pearson’s test) reveals exact drought specificity. (C) hierarchical clustering analysis revealed that the transcriptomes were grouped mainly according to water deficit intensity. RNA-seq libraries abbreviations for *S. chilense*, CL-A to CL-D: leaves in control condition, CSA-E to CSA-H: shoot apex in control condition, DL-I to DL-L: leave in drought condition, DSA-M to DSA-P: shoot apex in drought condition; SP-C1 to SP-C4: *S. pennellii* transcriptomes in control condition, SP-D1 to SP-D2: *S. pennellii* transcriptomes in drought condition; SL-C: *S. lycopersicum* transcriptome in control condition, SL-D: *S. lycopersicum* transcriptome in drought condition. Color scale indicates correlation coefficients from high values in red to low values in white.

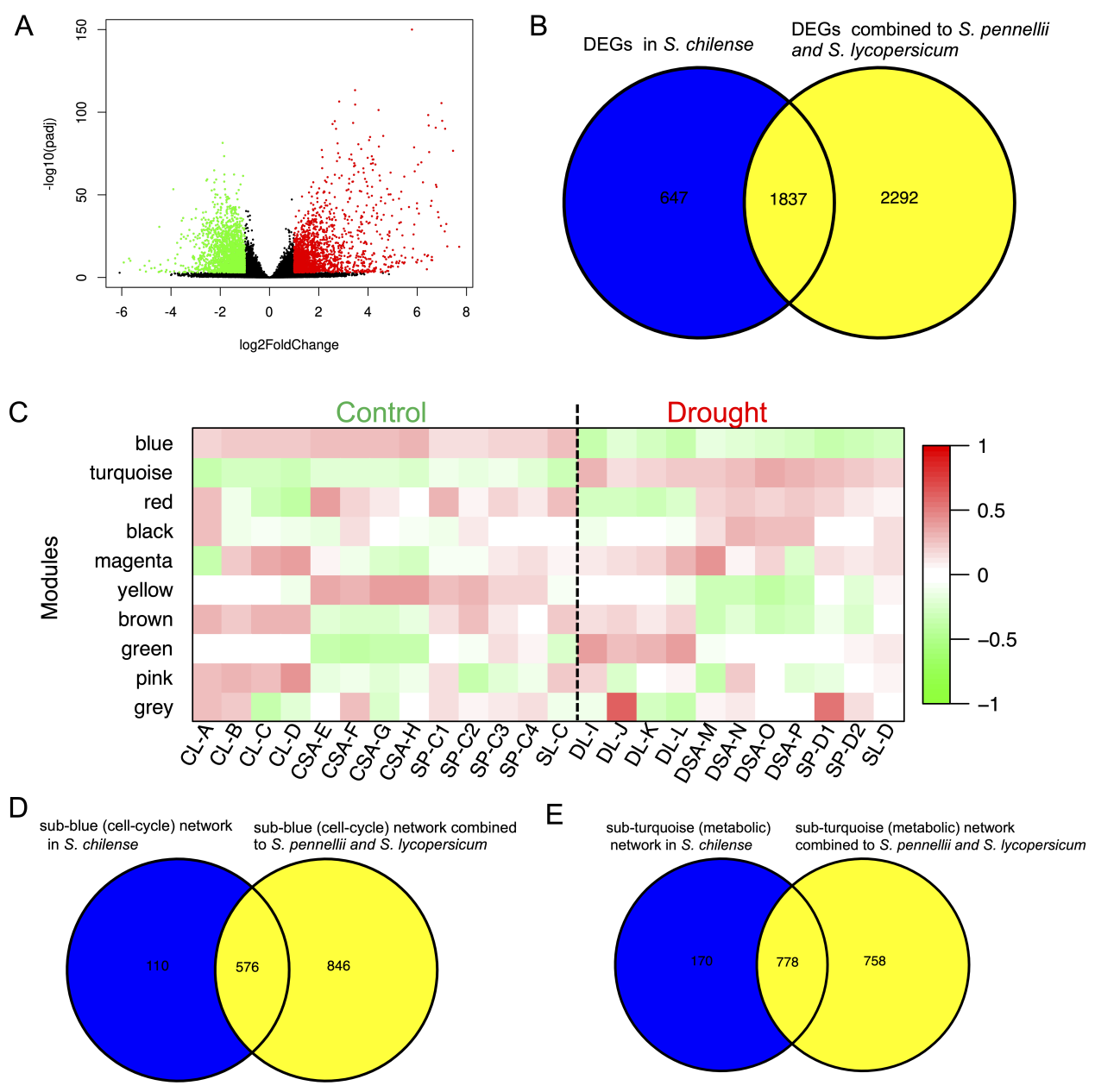

**Figure S4**. Identification of drought-response networks combined to transcriptomes of *S. pennellii* and *S. lycopersicum*. (A) Differentially Expressed Genes (DEGs) identified based on all control and drought transcriptomes. (B) Venn plot show the overlapped genes between DEG sets from only *S.chilense* and combining to *S. pennellii* and *S. lycopersicum*. (C) The correlation (Kendall's test) between samples expression patterns for the ten modules based on transcriptomes of *S. chilense*, of *S. pennellii* and *S. lycopersicum*. Color scale indicates correlation coefficients from high positive coefficient in red to high negative coefficient in green. No correlation is indicated in white. (D) and (E) The overlaps between networks based on only *S. chilense* and combining to *S. pennellii* and *S. lycopersicum* in sub-blue and sub-turquoise, respectively.

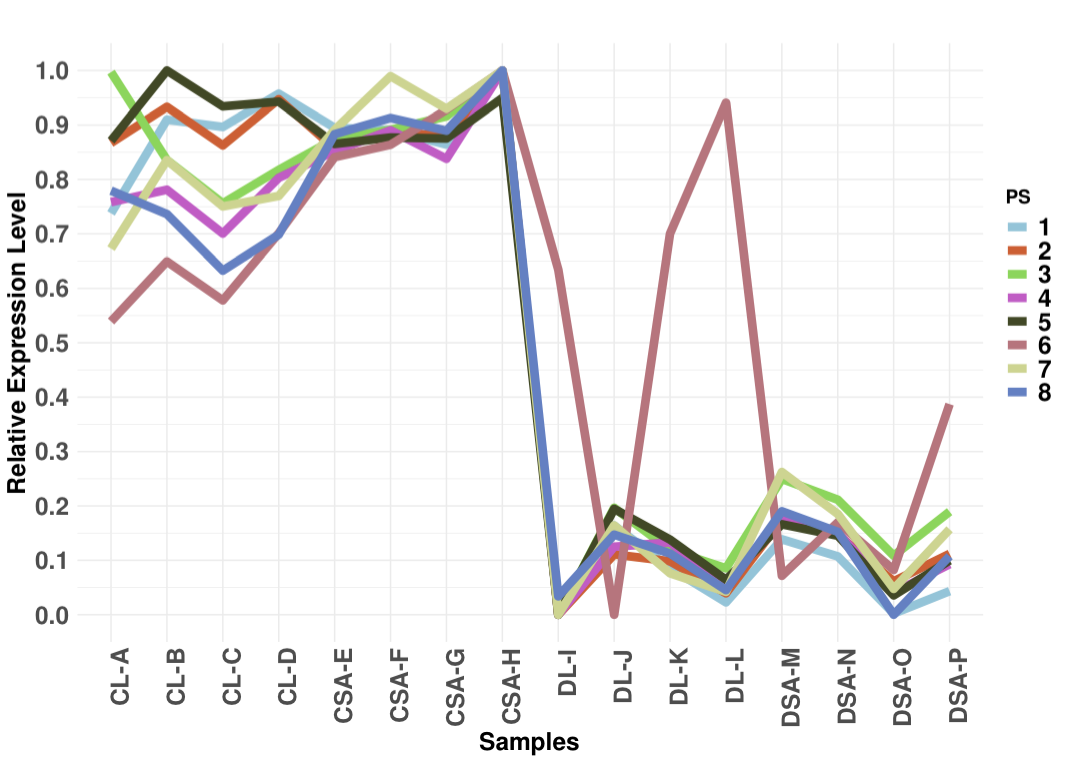

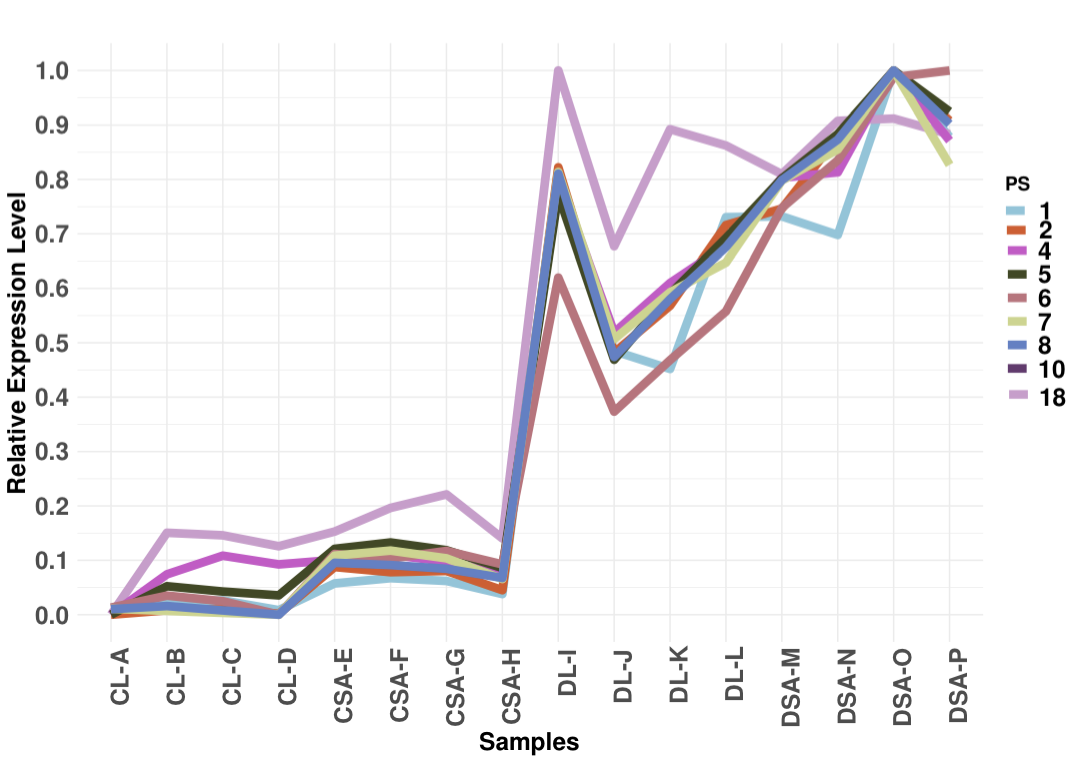
**Figure S5**. Relative expression level of genes in different Phylostrata (PS) in (A) cell-cycle and (B) metabolic networks depending on condition and tissue: CL-A to CL-D: leaves in control condition, CSA-E to CSA-H: shoot apex in control condition, DL-I to DL-L: leaves in drought condition, and DSA-M to DSA-P: shoot apex in drought condition.

A

BB

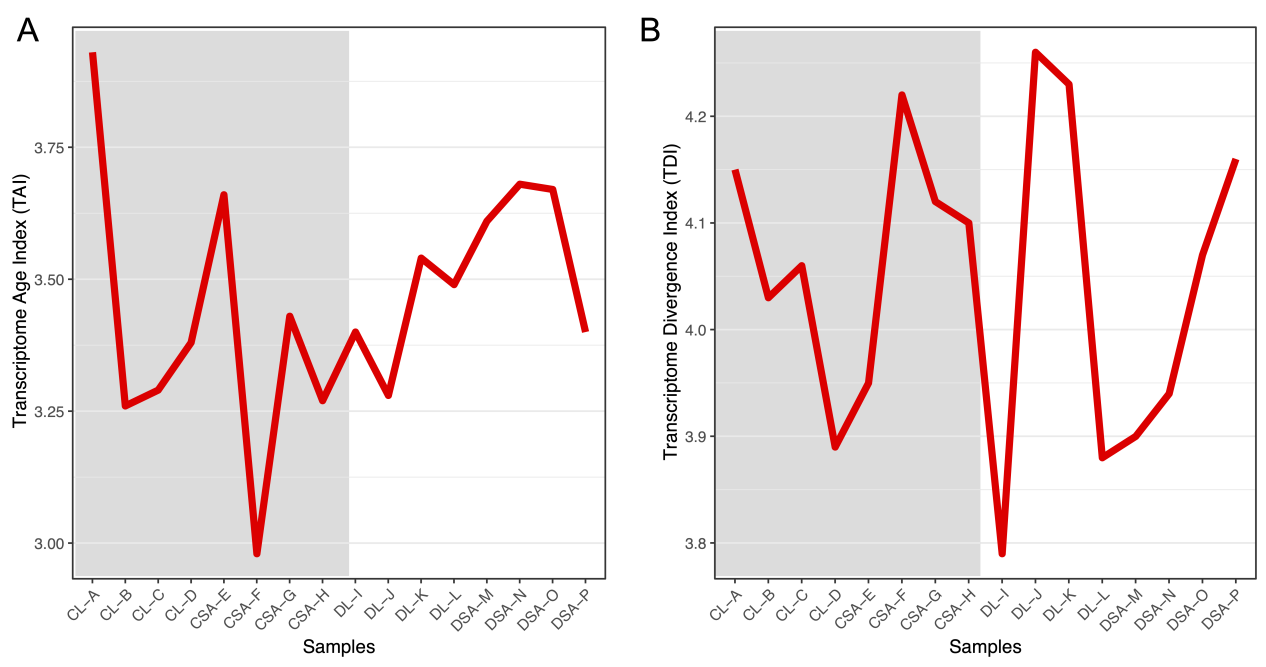

**Figure S6**. Transcriptome Age Index (TAI) and Transcriptome Divergence Index (TDI) in control (grey background) and drought (white background) based on 1,000 randomly selected genes and these genes are not in gene sets of differential expression gene, cell-cycle network and metabolic network.

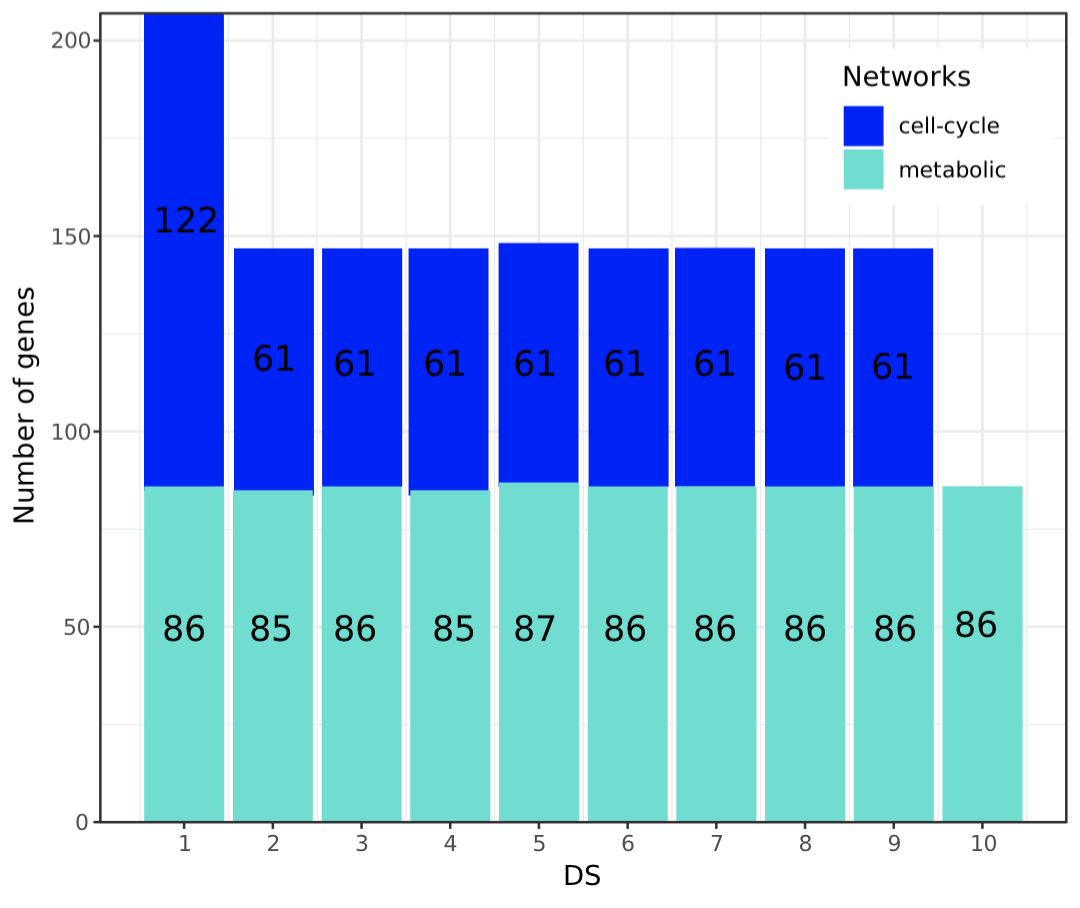

**Figure S7**. Number of genes in each divergence strata (DS) for two networks based on Ka/Ks.

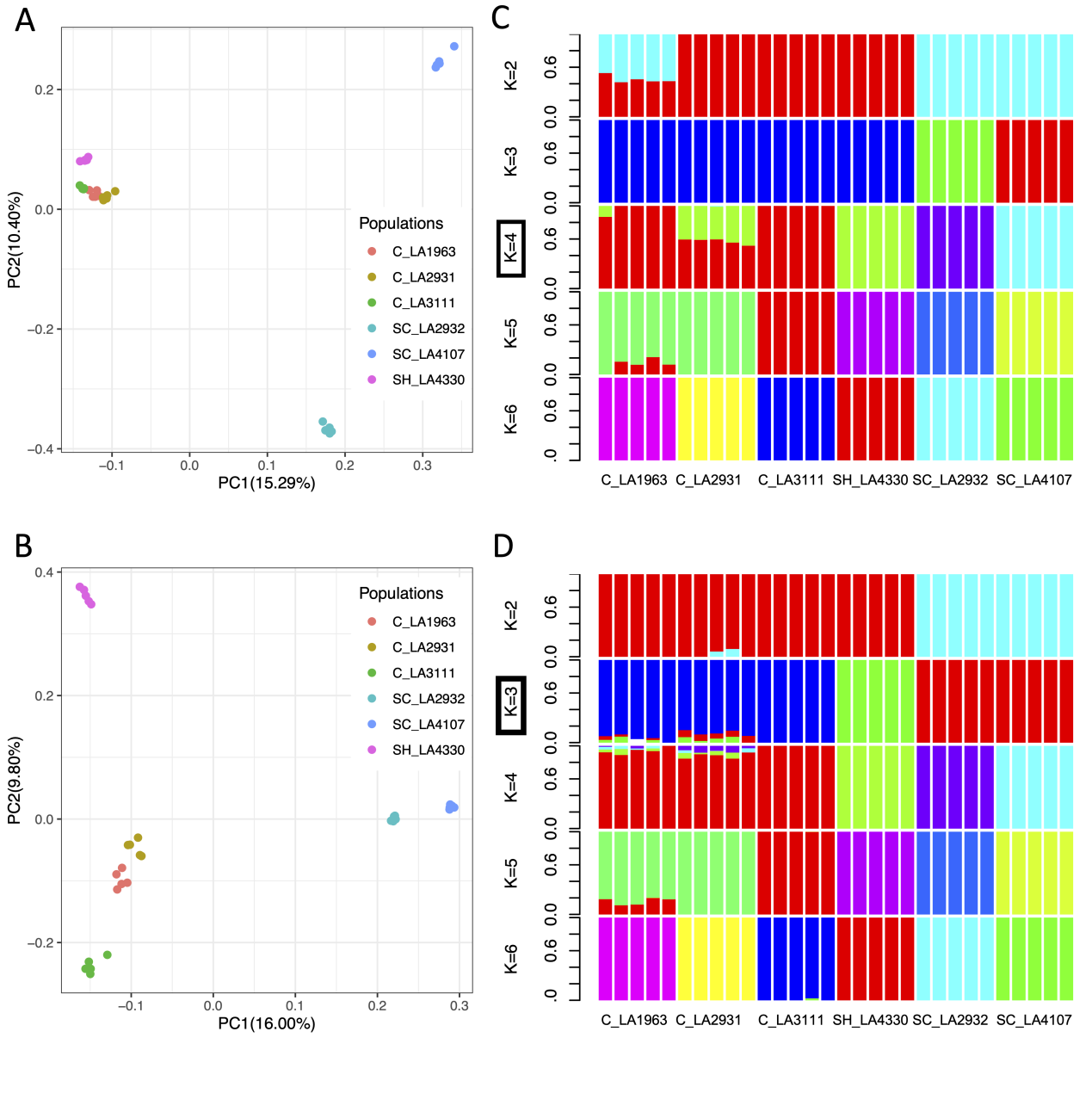

**Figure S8**. Population structure analysis using SNPs from 30 full-genome sequencing data. (A) PCA plot based on all SNPs in the genome. (B) PCA plot based on SNPs in regions of drought-responsive genes. (C) Structure analysis assuming K = [2 - 6] subpopulations based on all SNPs in the genome. The black box denotes the best K value. (D) Structure analysis based on SNPs in regions of drought-responsive genes.

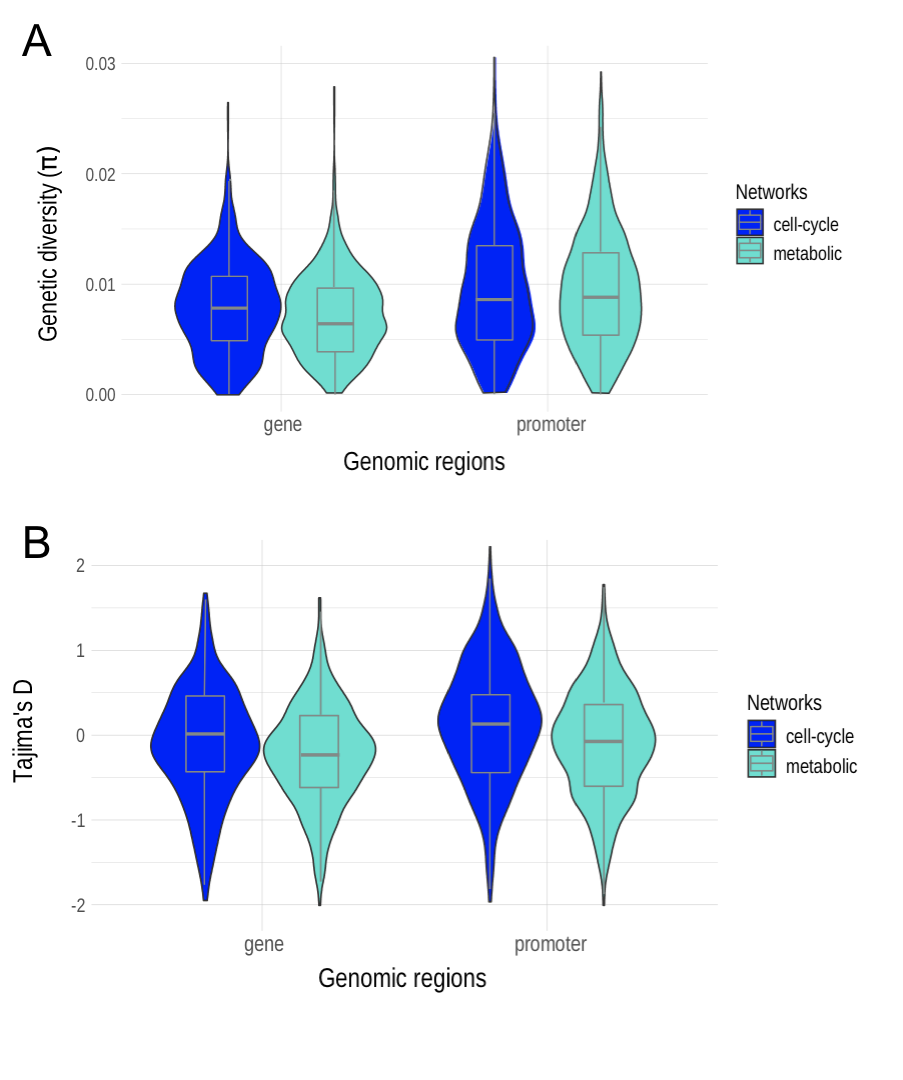
**Figure S9**. Statistics of population genetics at drought-responsive genes in the two chosen networks. (A) Nucleotide diversity (π) of gene (coding region) and promoter regions. (B) Tajima’s D of gene (coding region) and promoter regions. The values are computed from the 5 full genomes for the population LA1963.

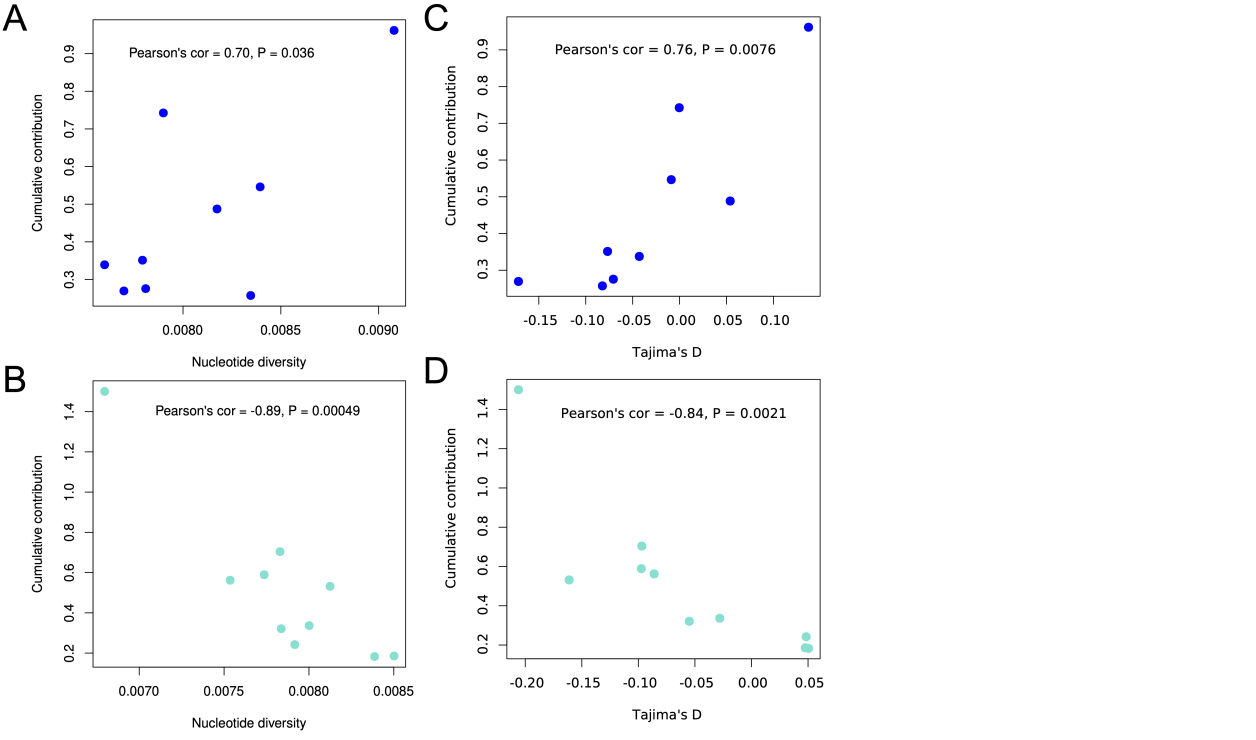

**Figure S10.** Correlation between nucleotide diversity (π) and contribution of each divergence strata (DS) in (A) cell-cycle and (B) metabolic networks. Correlation between Tajima’s D and contribution of each divergence strata (DS) in (C) cell-cycle and (D) metabolic networks.

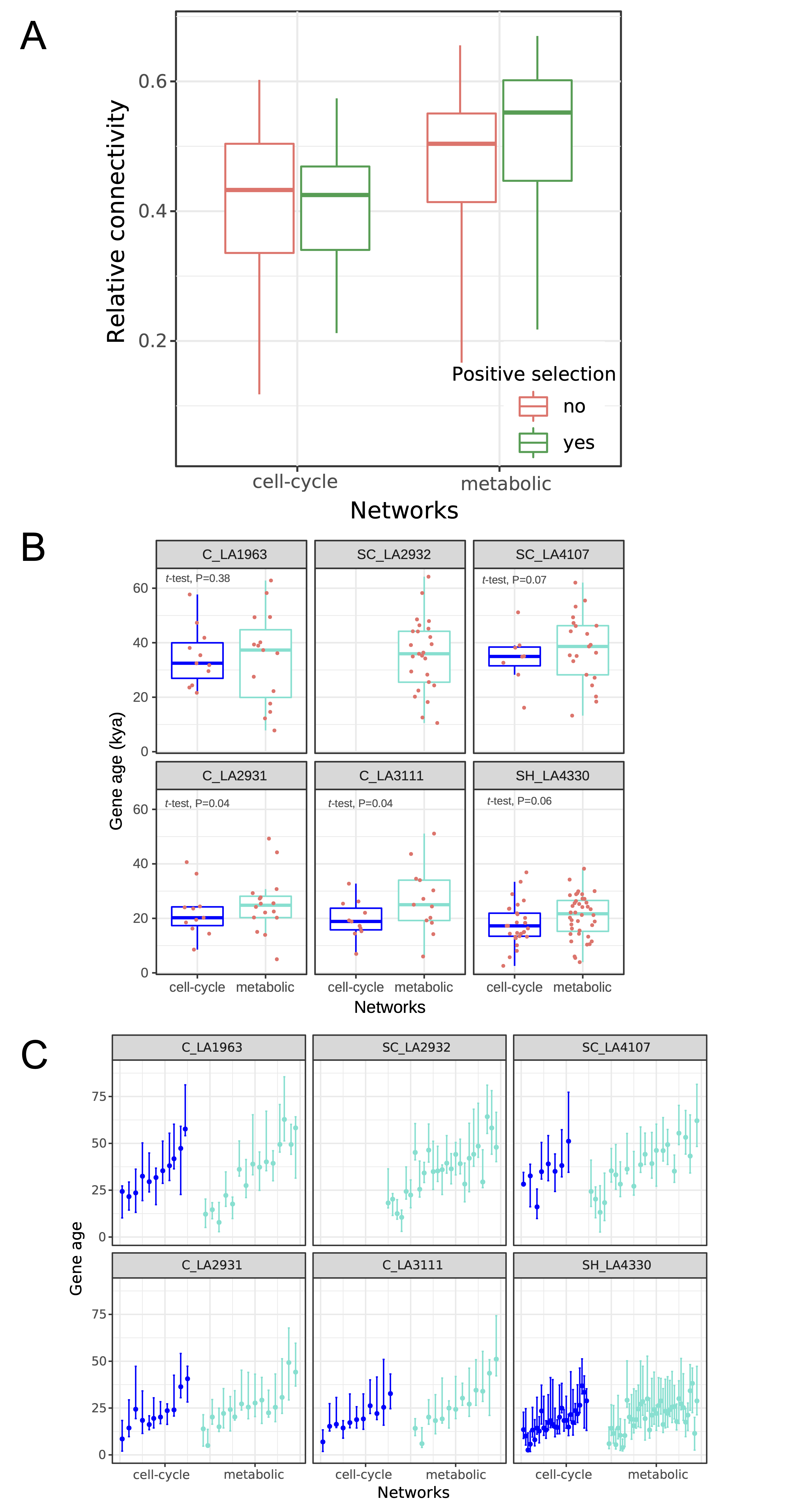

**Figure S11.** The connectivity and age of selective sweeps at genes of the two networks. (A) The relative connectivity between positive selected genes and other genes in the two drought-responsive networks. (B) Difference of the selective sweep age between positively selected genes of the cell-cycle and metabolic networks in six populations. (C) The distribution of the selective sweep age in the two networks and for each six population with uncertainty estimates from McSwan (Tournebize *et al*. 2019, as used in Wei *et al*. 2023).

**Table S1**. Shared genes between gene sets of DEGs and modules.

| Gene set of DEGs | red (474) | blue (3,852) | yellow (1,557) | turquoise (5364) | brown (3,666) | green (774) | black (183) |
| --- | --- | --- | --- | --- | --- | --- | --- |
| Drought vs Control (all samples: 4,012) | 6 | 1,812 | 113 | 1,660 | 90 | 0 | 0 |
| Drought vs Control (leaf: 3,692) | 49 | 1,760 | 112 | 1,440 | 200 | 16 | 0 |
| Drought vs Control (shoot apex: 3,367) | 22 | 1,487 | 150 | 1,457 | 11 | 0 | 5 |
| Drought vs Control (total: 4,905) | 56 | 2,101 | 192 | 1,947 | 225 | 16 | 5 |
| Drought vs Control (overlaps: 2484) | 4 | 1,223 | 70 | 1,079 | 67 | 0 | 0 |

Note: we do not check grey module because genes in grey module do not show any co-expression with other genes.

All samples denote that DEGs are identified between 8 control samples and 8 drought samples, leaf and shoot apex denote that DEGs are identified between 4 control samples and 4 drought samples for leaf and shoot apex, respectively. Total denote all DEGs identified in three comparison groups, overlaps denote shared DEGs in three comparison groups.

**Table S2**. The number of TFs and its TFBSs in two networks

| TF family | sub-blue | | sub-turquoise | |
| --- | --- | --- | --- | --- |
|  | TF | TFBS | TF | TFBS |
| BBR-BPC |  |  | 2 | 45 |
| bHLH | 5 | 1 | 9 | 3 |
| bZIP |  |  | 8 | 15 |
| C2H2 | 1 | 28 |  |  |
| Dof | 1 | 90 | 2 | 361 |
| E2F/DP | 2 | 3 |  |  |
| ERF | 2 | 40 | 6 | 158 |
| FAR1 |  |  | 1 | 1 |
| GATA | 1 | 11 | 1 | 30 |
| GRAS | 1 | 39 | 3 | 147 |
| HSF |  |  | 8 | 1 |
| LBD |  |  | 1 | 23 |
| MIKC_MADS |  |  | 1 | 32 |
| MYB | 3 | 54 | 6 | 37 |
| MYB_related | 1 | 5 |  |  |
| NAC | 1 | 2 |  |  |
| TALE | 1 | 410 | 3 | 33 |
| TCP |  |  | 4 | 10 |
| Trihelix |  |  | 4 | 7 |

**Table S3**. Enrichment of biological process in two networks

| Terms of sub-blue | Number of genes | Terms of sub-turquoise | Number of genes |
| --- | --- | --- | --- |
| DNA replication initiation | 8 | response to heat | 23 |
| double-strand break repair via break-induced replication | 7 | response to water deprivation | 23 |
| ribosomal large subunit assembly | 10 | SCF-dependent proteasomal ubiquitin-dependent protein catabolic process | 11 |
| cell cycle | 16 | protein complex oligomerization | 8 |
| ribosomal large subunit biogenesis | 7 | response to hydrogen peroxide | 11 |
| cell division | 21 | peptidyl-tyrosine dephosphorylation | 8 |
| microtubule-based movement | 11 | cellular amino acid catabolic process | 6 |
| DNA replication | 11 | lipid storage | 6 |
| mitotic cell cycle | 6 | killing of cells of other organism | 6 |
| DNA-dependent DNA replication | 4 | response to salt stress | 21 |
| protein folding | 14 | galactose metabolic process | 5 |
| anaphase-promoting complex-dependent catabolic process | 4 | cinnamic acid biosynthetic process | 5 |
| spliceosomal snRNP assembly | 4 | aromatic amino acid family catabolic process | 5 |
| hydrotropism | 4 | erythrose 4-phosphate/phosphoenolpyruvate family amino acid catabolic process | 5 |
| RNA modification | 12 | positive regulation of transcription, DNA-templated | 16 |
| DNA repair | 13 | cuticle development | 4 |
| maturation of LSU-rRNA | 3 | L-phenylalanine metabolic process | 5 |
| embryo development ending in seed dormancy | 14 | cellular response to heat | 6 |
| mitochondrion organization | 4 | lipid droplet organization | 4 |
| rRNA metabolic process | 4 | negative regulation of abscisic acid-activated signaling pathway | 6 |
| ncRNA processing | 3 | cellular response to unfolded protein | 4 |
| ribosomal small subunit biogenesis | 3 | abscisic acid-activated signaling pathway | 11 |
| regulation of cyclin-dependent protein serine/threonine kinase activity | 5 | response to abscisic acid | 17 |
| negative regulation of translation | 4 | signal transduction | 14 |
| response to cadmium ion | 10 | protein folding | 15 |
| polar nucleus fusion | 3 | seed maturation | 4 |
| DNA recombination | 7 | translational initiation | 12 |
| cytokinesis by cell plate formation | 3 | ubiquitin-dependent protein catabolic process | 15 |
| rRNA processing | 8 | negative regulation of transcription, DNA-templated | 9 |
| pseudouridine synthesis | 4 | DNA methylation | 4 |
| ribosome biogenesis | 6 | response to freezing | 3 |
| cellular response to cold | 3 | negative regulation of endopeptidase activity | 8 |
| ribosomal small subunit assembly | 3 | arginine biosynthetic process | 3 |
| RNA biosynthetic process | 7 | stomatal movement | 3 |
| lipid metabolic process | 9 | protein dephosphorylation | 11 |
| fatty acid biosynthetic process | 6 | negative regulation of nucleic acid-templated transcription | 5 |
| meiotic cell cycle | 3 | lipid metabolic process | 11 |
| chromatin organization | 5 | protein polyubiquitination | 5 |
| floral organ abscission | 2 | regulation of transcription by RNA polymerase II | 16 |
| histone deacetylation | 2 | intracellular signal transduction | 10 |
| phosphate ion transmembrane transport | 2 | chromatin silencing | 3 |
| histone modification | 3 | negative regulation of seed germination | 3 |
| microtubule-based process | 4 | chaperone-mediated protein folding | 3 |
| DNA duplex unwinding | 12 | negative regulation of cellular macromolecule biosynthetic process | 5 |
| regulation of anthocyanin biosynthetic process | 2 | nuclear-transcribed mRNA catabolic process, nonsense-mediated decay | 3 |
| DNA endoreduplication | 2 | transcription elongation from RNA polymerase II promoter | 3 |
| cell redox homeostasis | 2 | jasmonic acid mediated signaling pathway | 4 |
| positive regulation of translational fidelity | 2 | response to ethylene | 5 |
| regulation of meristem development | 2 | regulation of abscisic acid-activated signaling pathway | 3 |
| response to brassinosteroid | 3 | rhythmic process | 3 |
| protein peptidyl-prolyl isomerization | 5 | regulation of secondary shoot formation | 3 |
| glutamine metabolic process | 3 | cold acclimation | 4 |
| negative regulation of cell population proliferation | 2 | protein refolding | 4 |
| cytoskeleton organization | 4 | protein sumoylation | 3 |
| nucleosome assembly | 5 | cellular response to cold | 3 |
| cytoplasmic microtubule organization | 2 | positive regulation of response to water deprivation | 3 |
|  |  | vegetative to reproductive phase transition of meristem | 5 |
|  |  | defense response to bacterium | 11 |
|  |  | phosphorelay signal transduction system | 6 |
|  |  | mRNA cis splicing, via spliceosome | 3 |
|  |  | root development | 8 |
|  |  | response to chitin | 7 |
|  |  | cellular response to hypoxia | 5 |
|  |  | cellular response to lipid | 3 |
|  |  | embryo sac development | 4 |
|  |  | gene silencing by RNA | 5 |
|  |  | regulation of circadian rhythm | 2 |
|  |  | positive regulation of translational elongation | 2 |
|  |  | positive regulation of translational termination | 2 |
|  |  | phosphate ion homeostasis | 2 |
|  |  | positive regulation of proteasomal ubiquitin-dependent protein catabolic process | 3 |

**Table S4.** Enrichment of cellular component in two networks

| Terms of sub-blue | Number of genes | Terms of sub-turquoise | Number of genes |
| --- | --- | --- | --- |
| cytosolic ribosome | 24 | SCF ubiquitin ligase complex | 11 |
| ribonucleoprotein complex | 38 | monolayer-surrounded lipid storage body | 6 |
| large ribosomal subunit | 15 | CCAAT-binding factor complex | 5 |
| nucleolus | 34 | protein-containing complex | 9 |
| microtubule | 18 | autophagosome | 3 |
| polysomal ribosome | 8 | cytoplasmic stress granule | 3 |
| small ribosomal subunit | 9 | nuclear speck | 4 |
| chromocenter | 5 | chromatin | 14 |
| MCM complex | 5 | lipid droplet | 3 |
| nucleosome | 11 | secretory vesicle | 4 |
| intracellular organelle lumen | 22 | vacuole | 16 |
| organelle lumen | 22 | P-body | 4 |
| chloroplast thylakoid | 8 | endoplasmic reticulum lumen | 3 |
| small-subunit processome | 7 | mitochondrial respiratory chain complex I | 2 |
| mitochondrial matrix | 8 | intrinsic component of membrane | 2 |
| preribosome, large subunit precursor | 6 | vacuolar membrane | 11 |
| chromosome | 6 | plant-type cell wall | 5 |
| cytosolic small ribosomal subunit | 3 | ribonucleoprotein complex | 11 |
| kinesin complex | 4 | organelle envelope | 3 |
| phragmoplast | 6 | cell wall | 10 |
| plastid stroma | 8 | Golgi cisterna membrane | 1 |
| plastid | 15 | cytoskeleton | 3 |
| plastid envelope | 6 | large ribosomal subunit | 2 |
| cyclin-dependent protein kinase holoenzyme complex | 5 | spliceosomal complex | 4 |
| Cul4-RING E3 ubiquitin ligase complex | 4 | microbody | 4 |
| plastid thylakoid | 3 | peroxisome | 2 |
| chloroplast stroma | 15 | plastid | 9 |
| plant-type cell wall | 7 | chloroplast thylakoid | 2 |
| anaphase-promoting complex | 3 | plant-type vacuole | 2 |
| nuclear envelope | 3 | endoplasmic reticulum | 12 |
| cell plate | 3 | phragmoplast | 2 |
| U5 snRNP | 2 | chloroplast stroma | 10 |
| intracellular non-membrane-bounded organelle | 6 | Golgi trans cisterna | 1 |
| condensed chromosome, centromeric region | 2 | intracellular non-membrane-bounded organelle | 3 |
| 90S preribosome | 2 | plastid envelope | 2 |
| site of double-strand break | 2 | nucleosome | 4 |
| replication fork | 2 |  |  |
| spindle microtubule | 2 |  |  |

**Table S5**. Summary of population genetics statistics for the two networks

| Network | Tajima’s D | | | Nucleotide diversity (π) | | | Ka/Ks |
| --- | --- | --- | --- | --- | --- | --- | --- |
|  | genes | promoter | genes under positive selection | genes | promoter | genes under positive selection |  |
| cell-cycle | 0.00653±0.688 | 0.00936±0.679 | -0.00274±0.102 | 0.00786±0.00416 | 0.00976±0.00580 | 0.00548±0.00215 | 0.279±0.333 |
| metabolic | -0.0471±0.664 | -0.0201±0.675 | -0.0692±0.164 | 0.00724±0.00418 | 0.00973±0.00540 | 0.00507±0.00244 | 0.329±0.331 |

**Table S6**. The number of drought-responsive genes in the two networks which are found to be under positive selection in each population (Wei *et al*. 2023)

| Populations | cell-cycle | metabolic |
| --- | --- | --- |
| C_LA1963 | 11 | 15 |
| C_LA2931 | 11 | 16 |
| SC_LA2932 | 0 | 25 |
| C_LA3111 | 11 | 13 |
| SC_LA4107 | 8 | 21 |
| SH_LA4330 | 33 | 36 |

Note: the numbers outside brackets represent the number of drought-responsive genes under positive selection in our previous study based on reference genome of *S. pennellii* (Wei *et al*. 2023), the numbers in brackets denote shared drought-responsive genes between genome scans based on reference genomes of *S. chilense* and *S. pennellii*.

**Table S7**. The mean connectivity and age of the selective sweep for genes in the two networks

| Population | Cell-cycle | |  | Metabolic | |
| --- | --- | --- | --- | --- | --- |
|  | connectivity | gene age* |  | connectivity | gene age* |
| C_LA1963 | 0.43 ± 0.04 | 34,890 ±10,918 |  | 0.50 ± 0.13 | 35,258 ± 16,865 |
| C_LA2931 | 0.45 ± 0.05 | 22,396 ± 9,267 |  | 0.49 ± 0.13 | 25,175 ± 10,653 |
| SC_LA2932 | - | - |  | 0.57 ± 0.06 | 35,522 ± 13,339 |
| C_LA3111 | 0.43 ± 0.09 | 19,525 ± 6,917 |  | 0.52 ± 0.093 | 26,773 ± 12,147 |
| SC_LA4107 | 0.39 ± 0.07 | 34,421 ± 9,938 |  | 0.52 ± 0.09 | 37,946 ± 12,817 |
| SH_LA4330 | 0.47 ± 0.09 | 17,925 ± 7,634 |  | 0.51 ± 0.10 | 20,847 ± 7,984 |

*Gene age refers to the age of sweep region where the gene is located.
